# Aβ(1-42) tetramer and octamer structures reveal edge pores as a mechanism for membrane damage

**DOI:** 10.1101/759472

**Authors:** Sonia Ciudad, Eduard Puig, Thomas Botzanowski, Moeen Meigooni, Andres S. Arango, Jimmy Do, Maxim Mayzel, Mariam Bayoumi, Stéphane Chaignepain, Giovanni Maglia, Sarah Cianferani, Vladislav Orekhov, Emad Tajkhorshid, Benjamin Bardiaux, Natàlia Carulla

## Abstract

The formation of amyloid-beta (Aβ) oligomer pores in the membrane of neurons has been proposed as the means to explain neurotoxicity in Alzheimer’s disease (AD). It is therefore critical to characterize Aβ oligomer samples in membrane-mimicking environments. Here we present the first three-dimensional structure of an Aβ oligomer formed in dodecyl phosphocholine (DPC) micelles, namely an Aβ(1-42) tetramer. It comprises a β-sheet core made of six β-strands, connected by only two β-turns. The two faces of the β-sheet core are hydrophobic and surrounded by the membrane-mimicking environment. In contrast, the edges of the core are hydrophilic and are solvent-exposed. By increasing the concentration of Aβ(1-42), we prepared a sample enriched in Aβ(1-42) octamers, formed by two Aβ(1-42) tetramers facing each other forming a β-sandwich structure. Notably, samples enriched in Aβ(1-42) tetramers and octamers are both active in lipid bilayers and exhibit the same types of pore-like behaviour, but they show different occurrence rates. Remarkably, molecular dynamics simulations showed a new mechanism of membrane disruption in which water and ion permeation occurred through lipid-stabilized pores mediated by the hydrophilic residues located on the core β-sheets edges of the Aβ(1-42) tetramers and octamers.

## Introduction

Substantial genetic evidence links the amyloid-β peptide (Aβ) to Alzheimer’s disease (AD) (*1*). However, there is great controversy in establishing the exact Aβ form responsible for neurotoxicity. Aβ is obtained from a membrane protein, the amyloid precursor protein (APP), through the sequential cleavage of β- and γ-secretase (*2*). It is generally considered that, upon APP cleavage, Aβ is released to the extracellular environment. Due to its hydrophobic nature, Aβ then aggregates into multiple species, commonly referred to as soluble Aβ oligomers (*3*), which eventually evolve into Aβ fibrils (*4*-*7*), the main component of amyloid plaques. However, by focusing exclusively on Aβ aggregation in solution, the membrane is overlooked. Indeed, a large number of studies have shown that the interaction of Aβ with the membrane results in the formation of membrane-associated oligomers, whose formation is considered to be directly responsible for compromising neuronal membrane integrity (*8*-*12*).

In 2016, we reported conditions to prepare homogenous and stable Aβ oligomers in membrane-mimicking environments (*13*). We found that their formation was specific for Aβ(1-42)—the Aβ variant most strongly linked to AD—, that they adopted a specific β-sheet structure, which is preserved in a lipid environment provided by bicelles, and that they incorporated into membranes exhibiting various types of pore-like behaviour. Because of these properties, we named them β-barrel pore-forming Aβ(1-42) oligomers 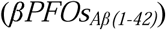. Here we present the atomic structures of 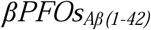 by NMR and MS and provide a mechanism for membrane disruption based on electrophysiology experiments and simulation studies in membranes.

### 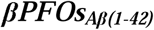 sample comprise Aβ(1-42) tetramers

To characterize 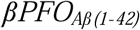, we prepared a selectively labeled ^2^H,^15^N,^13^C 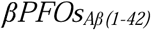 sample in dodecylphosphocholine (DPC) micelles at pH 9.0—conditions under which the sample was most stable (*13*)—and used high field nuclear magnetic resonance (NMR) triple-resonance TROSY-type experiments to obtain sequence-specific resonance assignments (fig. S1 and fig. S2). Peak assignment allowed us determining that Aβ(1-42) residues were observed in duplicate in the 2D [^1^H,^15^N]-TROSY spectrum (Fig. 1A), which suggested that the sample comprised two distinct Aβ(1-42) subunits. To highlight the detection of two Aβ(1-42) subunits in the sample, residues belonging to each of them were identified as either “red” or “green”. Next, we used the Cα and Cβ chemical shifts to derive the three-residue averaged (ΔCα-ΔCβ) secondary chemical shifts to thus determine the presence of secondary structure elements in each Aβ(1-42) subunit (Fig. 1B). This analysis revealed that the red Aβ(1-42) subunit contributed two β-strands, β1 and β2, to the oligomer structure. These strands extended, respectively, from G9 to A21 and from G29 to V40. Instead, in the green Aβ(1-42) subunit, residues L17 to F20 exhibited α-helical propensity, while residues G29 to I41 adopted a β-strand conformation, referred to as α1 and β3, respectively. To finalize assignments, the connectivity between β1 and β2, and α1 and β3 secondary structural elements was established using mixtures of Aβ(1-42) and Aβ(17-42) with distinct isotope labels (fig. S3).

**Fig. 1.**
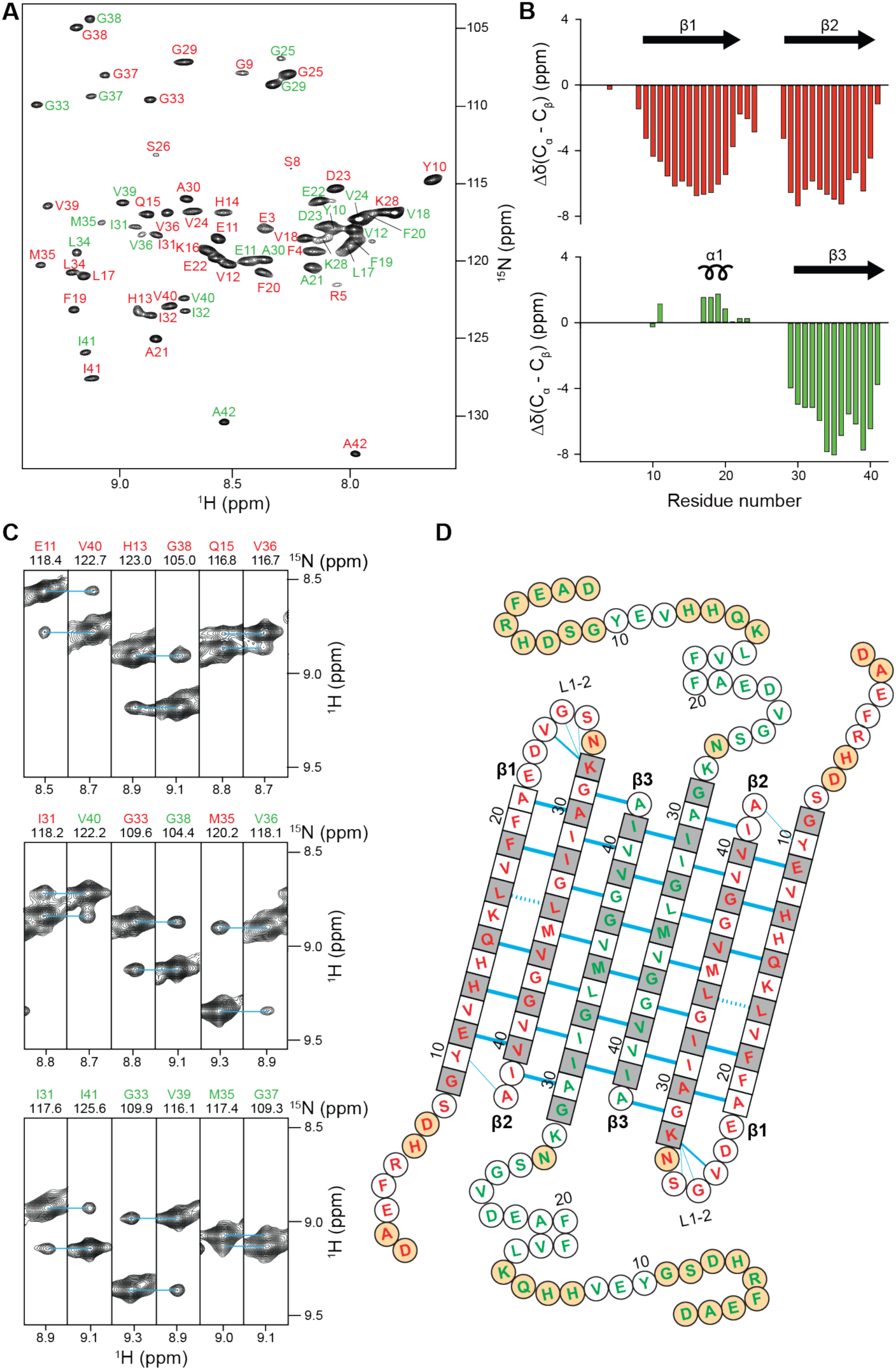
Architecture of the Aβ(1-42) tetramer. (**A**) Amide resonance assignments of the Aβ(1-42) tetramer. Two Aβ(1-42) subunits are detected and residues belonging to each of them are labeled in either red or green. (**B**) Three-bond averaged secondary chemical shifts versus residue number for the red (top) and the green (bottom) Aβ(1-42) subunits. Secondary structural elements derived from chemical shift indices are shown at the top with its corresponding number. Arrows indicate β-strands and helical symbols helices. (**C**) Strips from a 3D NH-NH NOESY spectrum defining long-range intra-monomer interactions between the red Aβ(1-42) subunit, long-range inter-monomer interactions between the red and the green Aβ(1-42) subunits, and long-range inter-dimer interactions between the two green Aβ(1-42) subunits. (**D**) The amino acid sequence of the Aβ(1-42) tetramer is arranged on the basis of the secondary and tertiary structure. Amino acids in square denote β-sheet secondary structure as identified by secondary chemical shifts; all other amino acids are in circles. Blue lines denote experimentally observed NOE contacts between two amide protons. Bold lines indicate strong NOEs typically observed between hydrogen-bonded residues in β-sheets. Dashed lines show probable contacts between protons with degenerate ^1^H chemical shifts. The side chains of white and gray residues point towards distinct sides of the β-sheet plane, respectively. Orange circles correspond to residues that could not be assigned.

Next, we used nuclear Overhauser effect spectroscopy (NOESY) to obtain long-range structural information. From the cross-peaks observed in the 3D NH-NH NOESY experiment, we identified 8 NOEs between β1 and β2 strands of the red Aβ(1-42) subunit and 7 NOEs between β2 strand of the red Aβ(1-42) subunit and the β3 strand of the green Aβ(1-42) subunit (Fig. 1C). The observation of intra- and inter-subunit NOEs allowed us to establish the topology of an asymmetric dimer unit and to confirm that all the peaks detected in the 2D [^1^H,^15^N]-TROSY spectrum belonged to the same oligomer. Moreover, we also detected three NOEs involving residues of the β3 strand (Fig. 1C), which could be explained only if two asymmetric dimer units interacted through β3 to form a tetramer in an antiparallel manner. All together, these NOEs allowed us to establish the complete topology of a six-stranded Aβ(1-42) tetramer unit (Fig. 1D). Moreover, since we did not detect any NOEs for the amide protons of β1 residues pointing outward of the β-sheet core (*i.e.,* Y10, V12, H14, K16, V18, and F20), we inferred that the signals detected by NMR corresponded to an Aβ(1-42) tetramer. To further validate the tetramer topology, we prepared specifically isotope-labeled samples and assigned the methyl groups of Ala, Ile, Leu, and Val (AILV) residues (fig. S4). We then acquired 3D NH-CH_3_ NOESY and 3D CH_3_-CH_3_ NOESY spectra and obtained a network of 87 NH-CH_3_ and 25 CH_3_-CH_3_ NOEs consistent with the topology of the tetrameric unit (fig. S5).

NMR NOESY-type experiments allowed us to identify a network of more than 150 NOE contacts (table S1) which, together with backbone dihedral angle (fig. S6) and hydrogen-bond restraints, allowed us to define the structure of an Aβ(1-42) tetramer (Fig. 2A, fig. S7 and table S2). The tetramer comprised a β-sheet core made of six β-strands, connected by only two β-turns, leaving two short and two long N-termini flexible ends, the latter comprising α1. The root mean square deviation (RMSD) of the Aβ(1-42) tetramer ensemble was 0.77 and 1.34 A□ for the backbone and the heavy atoms of the six-stranded β-sheet core, respectively. Notably, all residues on both faces of the β-sheet core were hydrophobic except for three basic residues (*i.e.,* H13, H14, and K16) located in β1, at the edges of the β-sheet core (Fig. 2B). On the other hand, residues making the β-turns and the flexible N-termini ends were hydrophilic except for those comprising α1.

**Fig. 2.**
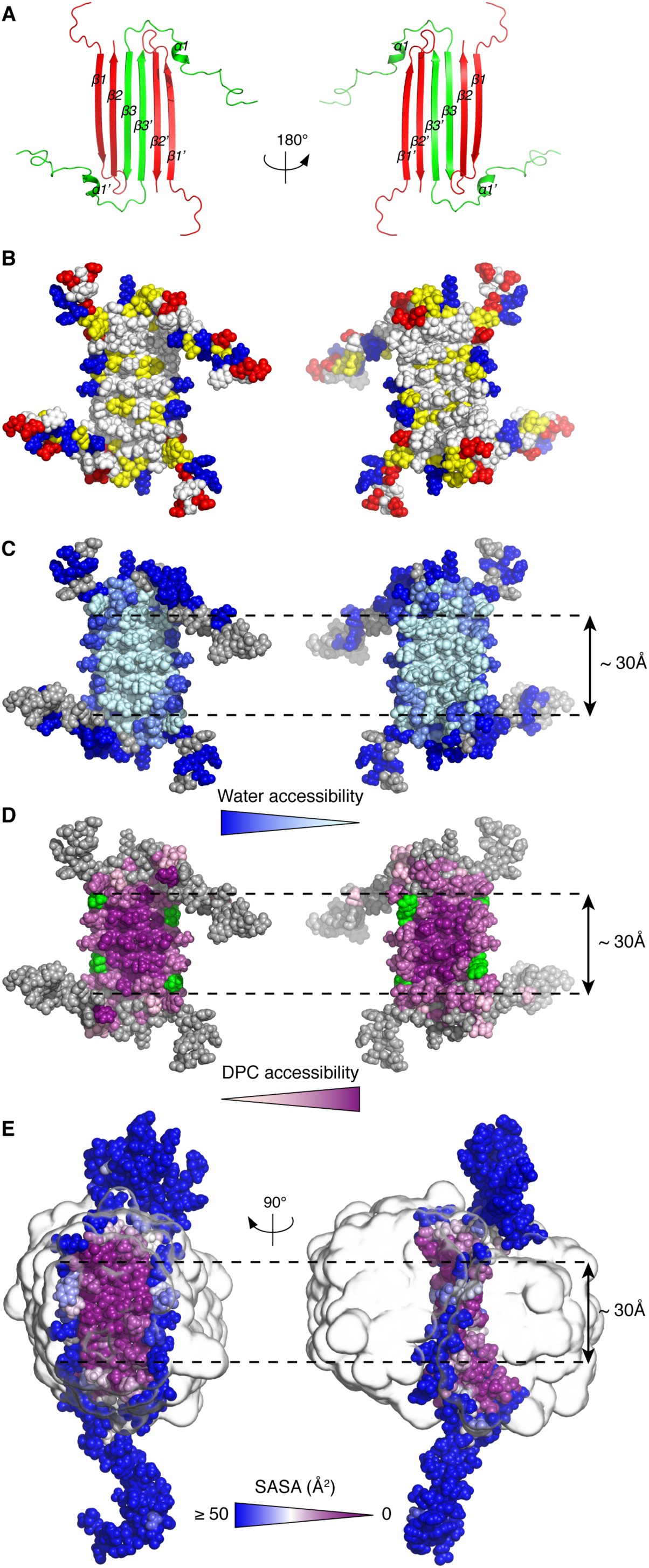
3D structure of the Aβ(1-42) tetramer prepared in DPC. (**A**) Ribbon diagram of the Aβ(1-42) tetramer structure. Aβ(1-42) subunits are colored either red or green to identify the asymmetric dimer unit that constitutes the building block of the Aβ(1-42) tetramer. (**B**) Distribution of hydrophobic and charged residues on the surface of the Aβ(1-42) tetramer. Hydrophobic residues are white, polar are yellow, and positively and negatively charged are red and blue, respectively. (**C**) Water accessibility of amide protons revealed through 2D [^1^H,^15^N]-HSQC spectra obtained at different pHs and through measurement of amide temperature coefficients. Solvent accessibility is linearly coded on the basis of the intensity of blue, with light blue corresponding to low water accessibility and dark blue corresponding to high water accessibility. Unassigned residues are shown in gray. (**D**) DPC accessibility of amide protons. The residues that showed NOEs between the backbone amide proton and the N-bound methyls of the choline head group of DPC are shown in green. The amide residues that showed paramagnetic enhancement, ε, upon addition of 16-DSA are shown in magenta. The ε values are linearly coded on the basis of the intensity of magenta, with light pink corresponding to ε = 0 and dark magenta corresponding to ε = ε_max_. (**E**) Solvent Accessible Surface Area (SASA, Å^2^) from MD simulations of the Aβ(1-42) tetramer in DPC. Detergent micelle is represented as a smoothed transparent surface. The figure was prepared with the program Pymol.

### Aβ(1-42) tetramer – DPC interaction

Having established the 3D structure and the physicochemical properties of the Aβ(1-42) tetramer, we examined how it interacted with the surrounding media, namely water and the DPC detergent molecules. 2D [^1^H,^15^N]-HSQC spectra were acquired at two pH 8.5 and 9.5 (fig. S8A,B). Residues belonging to the β-sheet core and some belonging to α1 were detected at both pHs, while some of the α1 residues and those corresponding to the β-turns and the N-termini ends were detected only when the spectrum was measured at the lowest pH. This observation thus suggests that residues comprising the β-turns and the N-termini ends exchanged faster with the solvent and were therefore more exposed than those making the β-sheet core and α1 (Fig. 2C). To establish whether the more protected β-sheet core residues exhibited distinct degrees of solvent protection, we determined their amide temsperature coefficients (Δδ/ΔT). Most of the NH amide protons of residues comprising β1, β2 and β3 were the most affected by temperature changes, which is consistent with these residues forming stable hydrogen bonds (*14*). In contrast, amide protons of β1 residues pointing out of the β-sheet core (*i.e.,* Y10, V12, H14, K16, V18, and F20) exhibited the lowest amide temperature coefficients, suggesting that these residues are the most water accessible of all residues comprising the β-sheet core (Fig. 2C). To characterize the interaction of the DPC molecules with the surface of the Aβ(1-42) tetramer, we acquired a 3D ^15^N-resolved [^1^H,^1^H]-NOESY spectrum of the Aβ(1-42) tetramer using a selectively ^13^C methyl-protonated AILV and otherwise uniformly ^2^H,^15^N Aβ(1-42) sample prepared using DPC at natural abundance. Analysis of this spectrum allowed us to identify two types of intermolecular interactions. First, we detected intermolecular NOEs between residues V12, L17, and L18, located in β1, and the N-bound methyl groups of the choline head group of DPC (Fig. 2D and fig. S9). Notably, this observation suggested that the detergent head group is bent towards the positively charged side chains (i.e., H13, H14, and K16) located at the hydrophilic edges of the β-sheet core in order to stabilize them. Second, we detected intermolecular NOEs between all amide protons comprising the β-sheet core and the hydrophobic tail of DPC, with the largest intensities for residues located at the center of the β-sheet core and decreasing toward its edges (Fig. 2D and fig. S9). These observations were confirmed using a paramagnetic labeled detergent, 16-doxyl stearic acid (16-DSA) (Fig. 2D and figs. S10, S11, and S12).

Finally, the interaction of the Aβ(1-42) tetramer with DPC micelles was further studied through molecular simulations using the SimShape approach (*16*). Over the course of a 1-ns simulation, the Aβ(1-42) tetramer was enveloped in a toroidal DPC micelle (fig. S13). Afterwards, the toroidal complex was equilibrated in explicit solvent for 60 ns. During this time, the hydrophobic terminal tail carbon of DPC was observed to interact predominantly with the two faces of the six-stranded β-sheet core, while transient contacts were also detected with the α1 region. Additionally, the DPC polar head was observed to interact with the hydrophilic edges of the six-stranded β-sheet region, which slowly became exposed to the solvent (Fig. 2E). Finally, these interactions were further validated by simulating the equilibrated protein-detergent complex in the absence of any external biasing forces (fig. S14). In summary, the experimental and the simulation results indicate that both faces of the central hydrophobic β-sheet core of the Aβ(1-42) tetramer were covered with a monolayer of DPC with α1 residues also interacting with the hydrophobic tail of DPC. In contrast, the rest of the residues, including the hydrophilic edges of the β-sheet core, were solvent-exposed and further stabilized by interactions with the polar head of DPC.

### Aβ(1-42) tetramers and octamers are present in 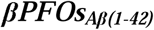 sample

Previous electrical recordings using planar lipid bilayers had revealed that the 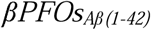 sample induced various types of pore-like behaviour (*13*). Having established the 3D structure of the Aβ(1-42) tetramer, it was difficult to envision how it could be directly responsible for pore formation. For this reason, we attempted to determine whether other oligomer stoichiometries, not detectable by NMR, were present in the 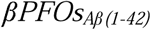 sample. To this end, we set to analyze the sample by means of size exclusion chromatography coupled to native ion mobility mass spectrometry (SEC/IM-MS) (*17*). This strategy presented a unique opportunity to establish the stoichiometry of the potentially distinct oligomer species present as a function of their elution through a SEC column. We had previously analyzed 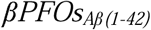 in a SEC column equilibrated in DPC and shown that the sample eluted as a major peak at 27.4 mL (Fig. 3A). However, to carry out SEC/IM-MS, a different detergent that would be compatible with MS analysis and would preserve oligomer stability was required (*15*). C8E5 was found to fulfill both requirements (Fig. 4A). MS analysis of the early eluting volume of the 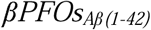 peak revealed charge states consistent with the presence of tetramers and octamers as the main species in the sample, respectively. Analysis of the late eluting volume showed an increase in the relative abundance of the charge states corresponding to tetramers relative to those assigned to octamers. Importantly, the use of IM prior to MS analysis allowed unambiguous assignment of the contribution of distinct oligomer stoichiometries to each charge state (fig. S15). This analysis led us to conclude that, although in agreement with NMR experiments, the stoichiometry of the major species present in the 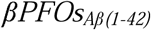 sample was Aβ(1-42) tetramers; Aβ(1-42) octamers were also present. In addition, since no charge states specific for other oligomer stoichiometries between tetramers and octamers were detected, these results suggested that tetramers were the building block for octamer formation. Notably, upon increasing activation conditions of the mass spectrometer, a population of tetramers broke into trimers and monomers (fig. S16). Instead, although the detection of octameric forms required slightly higher activation conditions than tetramers, once detected, octamers did not break at the maximum activation conditions afforded by the instrument. This result indicated that octamers were not derived from the forced co-habitation of two tetramers in a micelle but rather from specific interactions between the Aβ subunits composing it.

**Figure 3.**
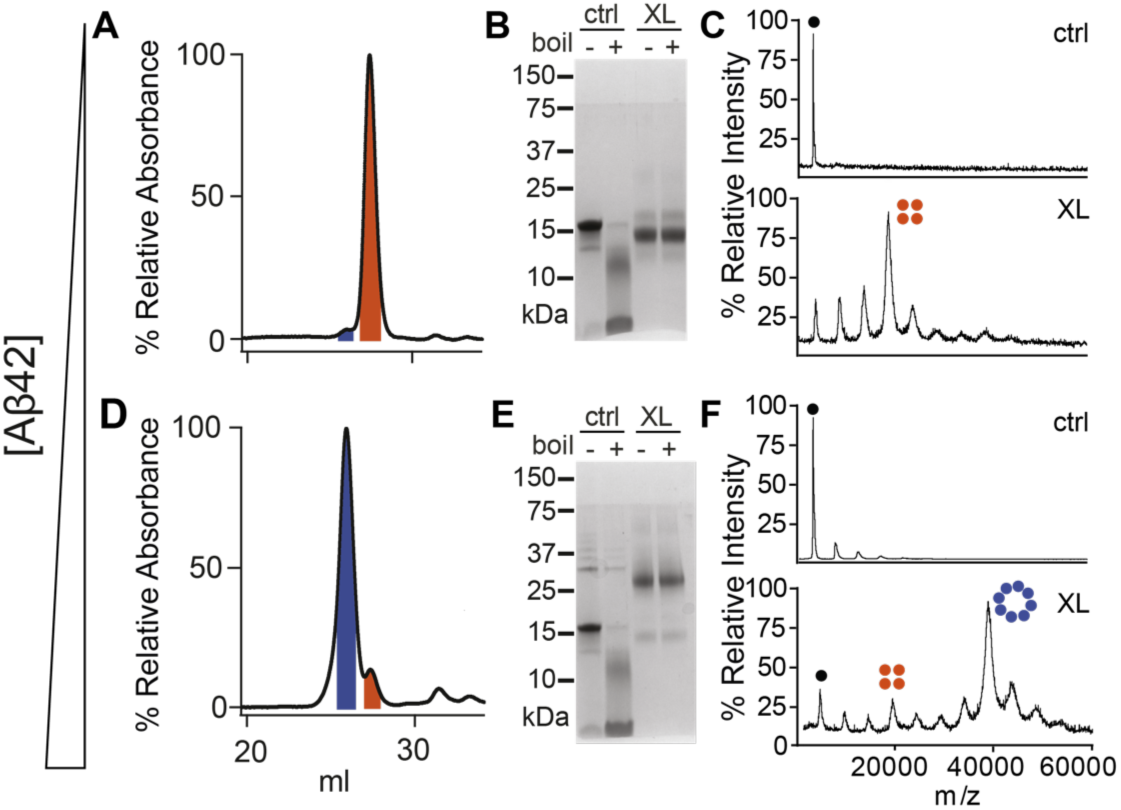
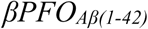 samples can be enriched in either tetramers or octamers. SEC of 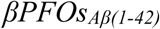 prepared at low (**A**) and high (**D**) Aβ(1-42) concentration in a column equilibrated in DPC. The peaks labeled in orange and blue are assigned, respectively, to Aβ(1-42) tetramers and octamers. SDS-PAGE analysis of 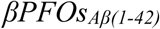 prepared at low (**B**) and high (**E**) Aβ(1-42) concentrations either not cross-linked (ctrl) or having been previously cross-linked (XL) and showing the effect of boiling (+) and non-boiling (-). MALDI-TOF analysis of 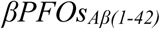 prepared at low (**C**) and high (**F**) Aβ(1-42) concentrations.

**Figure 4.**
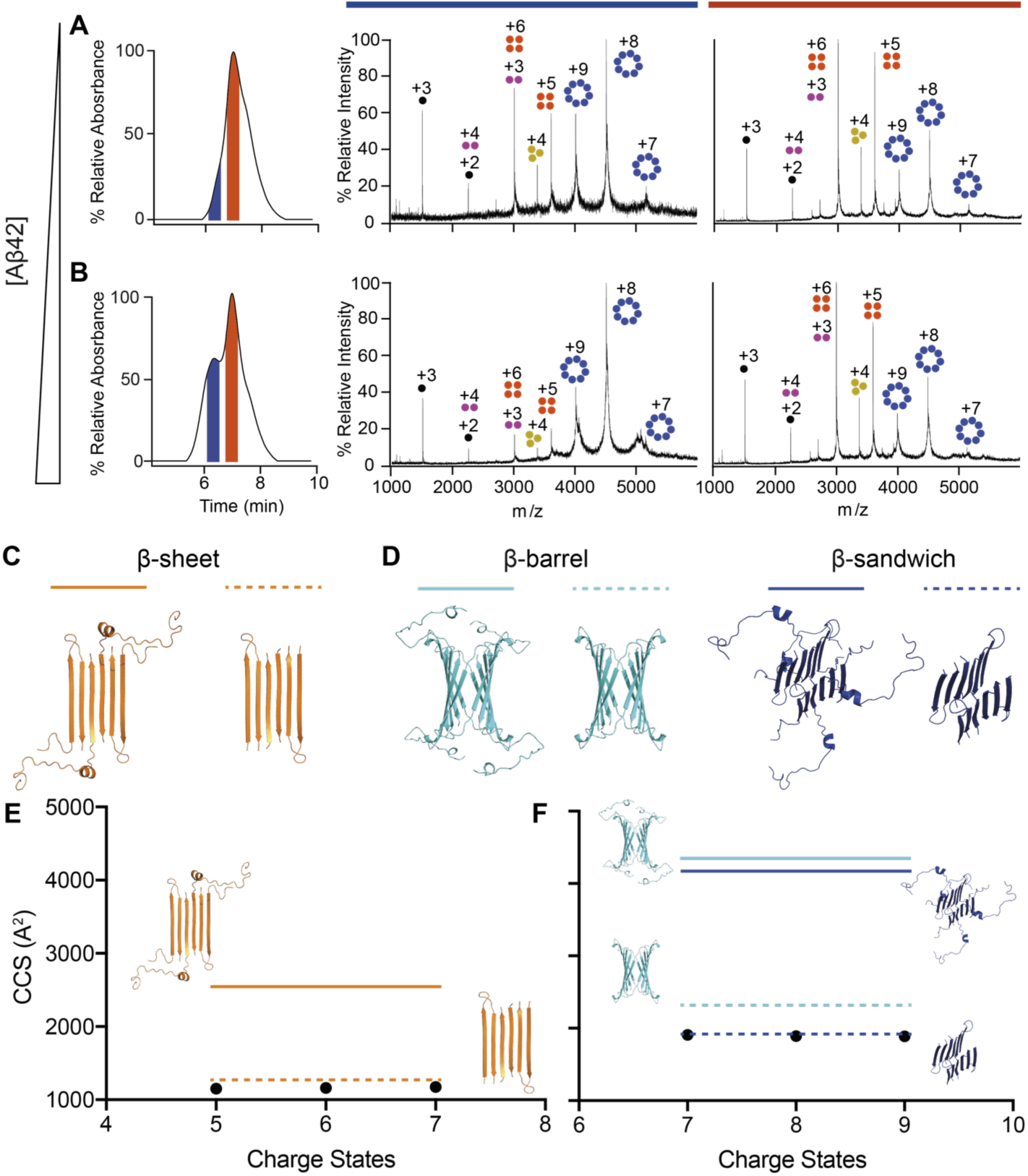
Aβ(1-42) octamers adopt a β-sandwich structure. SEC-MS analysis of 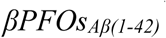 prepared at low (**A**) and high (**B**) Aβ(1-42) concentration. To couple SEC to MS analysis, the SEC column was equilibrated in C8E5. The mass spectra extracted from the blue and orange SEC peaks are shown, respectively, with a blue and orange line on top of them. The charge states corresponding to monomers, dimers, trimers, tetramers, and octamers are indicated with schematic drawings and labeled, respectively, in black, pink, yellow, orange and blue. (**C**) Aβ(1-42) tetramer structure derived from NMR restraints with (continuous line) and without (dashed line) the flexible N-terminal ends. **(D)** Octamer models based on the interaction of two Aβ(1-42) tetramers to form a β-barrel or a β-sandwich structure with (continuous line) and without (dashed line) the flexible N-terminal ends. (**E)** Experimental CCS of the tetramer (black dots) compared to the theoretical CCS of the Aβ(1-42) tetramer structure with and without flexible N-terminal ends. (**F)** Experimental CCS of the octamer (black dots) compared to the theoretical CCS of the four proposed octamer models.

### Preparation of a 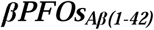 sample enriched in Aβ(1-42) octamers

To pursue the characterization of octameric species, we attempted to enrich our sample in this oligomer form. To this end, we maintained the concentration of DPC micelles constant and increased the concentration of Aβ(1-42) to mimic the consequences of an increase in the latter in the membrane (*12*). Thus, from this point, we worked with two 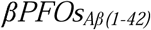 samples, one corresponding to the sample analyzed up to now and prepared at 150 μM of Aβ(1-42), referred to as 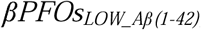, and one prepared at 450 μM Aβ(1-42), referred to as 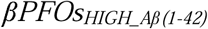. To establish whether 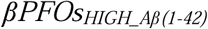 were enriched in octameric forms, we analyzed them by SEC using a column equilibrated in DPC (Fig. 3D). This analysis resulted in a major peak eluting 1.4 mL earlier than 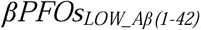as well as a small peak eluting at the same volume as the major peak detected for 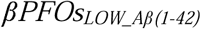. These findings indicated that working at high Aβ(1-42) concentration indeed led to the formation of a larger oligomer.

To study the stoichiometry of the oligomers present in the two samples, after preparing them in DPC micelles without any buffer exchange, we submitted them to chemical crosslinking. Given the abundance of basic and acid moieties in the flexible regions of the Aβ(1-42) tetramer structure derived by NMR (fig. S17), we decided to generate zero-length (ZL) cross-links between Lys and Asp or Glu residues using DMTMM as coupling reagent (*16*). As previously described, SDS-PAGE analysis of the non-cross-linked 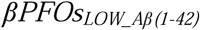 sample led, depending on whether the sample had been previously boiled or not, to either a 5 kDa band, corresponding to Aβ(1-42) monomers, or to a major band at 18 kDa, consistent with Aβ(1-42) tetramers (Fig. 3B) (*13*). In contrast, SDS-PAGE analysis of the cross-linked 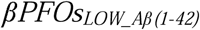 sample led to a major band at 14 kDa, regardless of whether it had been boiled previously. The increase in migration detected for the cross-linked samples is associated with protein compaction caused by crosslinking events (*17*). To further confirm the stoichiometry of the cross-linked bands established by SDS-PAGE, samples were analyzed by MALDI-TOF (Fig. 3C). MALDI ionization involves harsh conditions, which prevents preservation of the non-covalent interactions present in protein complexes. Therefore, as expected, the molecular weight of the sample analyzed by MALDI-TOF without being cross-linked led to the detection of a peak corresponding to the molecular mass of the monomer. Instead, analysis of the cross-linked 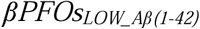 sample led to the detection of a major peak consistent with the mass of an Aβ(1-42) tetramer, thereby confirming the suitability of the ZL chemistry to efficiently cross-link the major species formed under this condition. Next, we applied the same cross-linking chemistry to the analysis of the 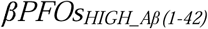 sample. SDS-PAGE analysis of the non-cross-linked samples led to the same bands obtained for 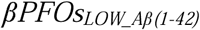as well as to a faint band at about 30 kDa, consistent with Aβ(1-42) octamers (Fig. 3E). Instead, SDS-PAGE analysis of the cross-linked 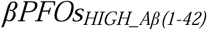 sample, both non-boiled and boiled, led to the detection of bands migrating at 28 kDa, consistent with Aβ(1-42) octamer formation. This result was further validated by MALDI-TOF analysis (Fig. 3F). All together, these results indicated that the 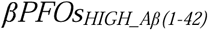 sample comprises mainly Aβ(1-42) octamers. Moreover, the observation that SDS-PAGE analysis of the non-cross-linked and non-boiled 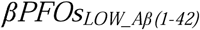 and 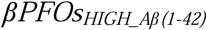 samples led to mainly the same Aβ(1-42) tetramer band points to Aβ(1-42) octamers being formed by two tetrameric building blocks whose stabilizing interactions are not preserved in the presence of SDS.

### Aβ(1-42) octamers adopt a β-sandwich structure with hydrophilic edges

Subsequently, we analyzed 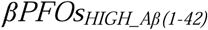 by SEC/IM-MS. Although C8E5, the detergent required for native MS analysis, did not completely stabilize the larger oligomer detected in a SEC column equilibrated in DPC (compare Fig. 3D to 4B), analysis of the early eluting peak, corresponding to the larger oligomeric species, led exclusively almost to three charge states assigned to Aβ(1-42) octamers (Fig. 4B, figs. S18 and S19). In summary, characterization of the 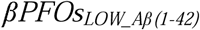 and 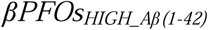 samples by SEC, cross-linking/MALDI-TOF and SEC/IM-MS revealed that the former was enriched in Aβ(1-42) tetramers and the latter in octamers.

To study the conformational state of the Aβ(1-42) octamers, we used IM-MS to derive their collision cross-sections (^TW^CCS_N2_) (Fig. 4C-F). The experimental ^TW^CCS_N2_ for the Aβ(1-42) tetramer was consistent only with the theoretical CCS obtained using the Aβ(1-42) tetramer structure determined by NMR, when the flexible loops were removed from the structure (Fig. 4C and 4E). This result indicated that these residues were partially collapsed in the gas phase, in line with observations made for other membrane proteins containing flexible loops (*18*). The experimental ^TW^CCS_N2_ for the Aβ(1-42) octamer was compared to two octamer models constructed using the 3D structure of the Aβ(1-42) tetramer as a building block (Fig. 4D). The first model was based on the association of two tetramers to form a loose β-barrel structure and the second one on the association of two tetramers in a β-sandwich structure. The experimental ^TW^CCS_N2_ for the Aβ(1-42) octamer was consistent with the theoretical CCS of the β-sandwich octamer when the flexible loops were removed (Fig. 4F). This result is indeed consistent with the physicochemical properties of the Aβ(1-42) tetramer as its two hydrophobic faces do not support its self-assembly in a β-barrel octamer structure with a central hydrophilic cavity. Instead, the Aβ(1-42) tetramer assembly in a β-sandwich octamer fully fulfils its physicochemical properties.

## Structures of the Aβ(1-42) tetramers and octamers reveal edge conductivity pores

Having obtained the means to prepare and characterize 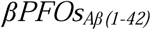 samples enriched in tetramers and octamers, we set to compare their activity in lipid bilayers by electrical recordings using planar lipid bilayers (fig. S20). The only difference between the two samples was found in the occurrence rate of the different pore-like behaviors with 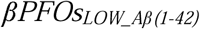enriched in tetramers, exhibiting fast and noisy transitions with undefined open pore conductance values for a higher number of times than 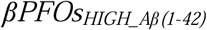, enriched in octamers, and the latter exhibiting a well-defined open pore with no current fluctuations for a higher number of times than the former. The similarity in ion conductance observed between the enriched Aβ(1-42) tetramer and octamer samples motivated the use of non-equilibrium molecular dynamics (MD) simulations to probe the mechanism of bilayer disruption at an atomistic scale. These simulations involved the application of an external electric field to observe ion conductance properties in 150 mM NaCl, 310K, at 100 mV. The presence of hydrophilic edges on both the membrane embedded Aβ(1-42) tetramer and octamer structures resulted in their unfavorable exposure to the hydrophobic lipid tails of the membrane. This situation led to lipid rearrangement, such that the head groups of the lipids reoriented to face the hydrophilic edges. This process led to the formation of lipid-stabilized pores, which stabilized the protein-lipid complex and allowed for water to permeate the membrane along the hydrophilic edges (Fig. 5). Contacts between protein and DPPC head group atoms were characterized for both tetramer and octamer systems (figs. S21 and S22). We observed a higher degree of water permeation and a greater solvent-accessible surface area in the octamer than in the tetramer (fig. S23). Although there were no ions that completely translocated across the membrane through the lipid-stabilized pore, ion permeation into the membrane space along the hydrated edges of the protein was observed during the simulations with applied electric field. We associate the formation of lipid-stabilized pores observed during the MD simulations with the mechanism of water and ion permeation observed experimentally through electrical recordings using planar lipid bilayers (fig. S20) and propose them to explain the neurotoxicity observed in AD through the disruption of cellular ionic homeostasis.

**Figure 5.**
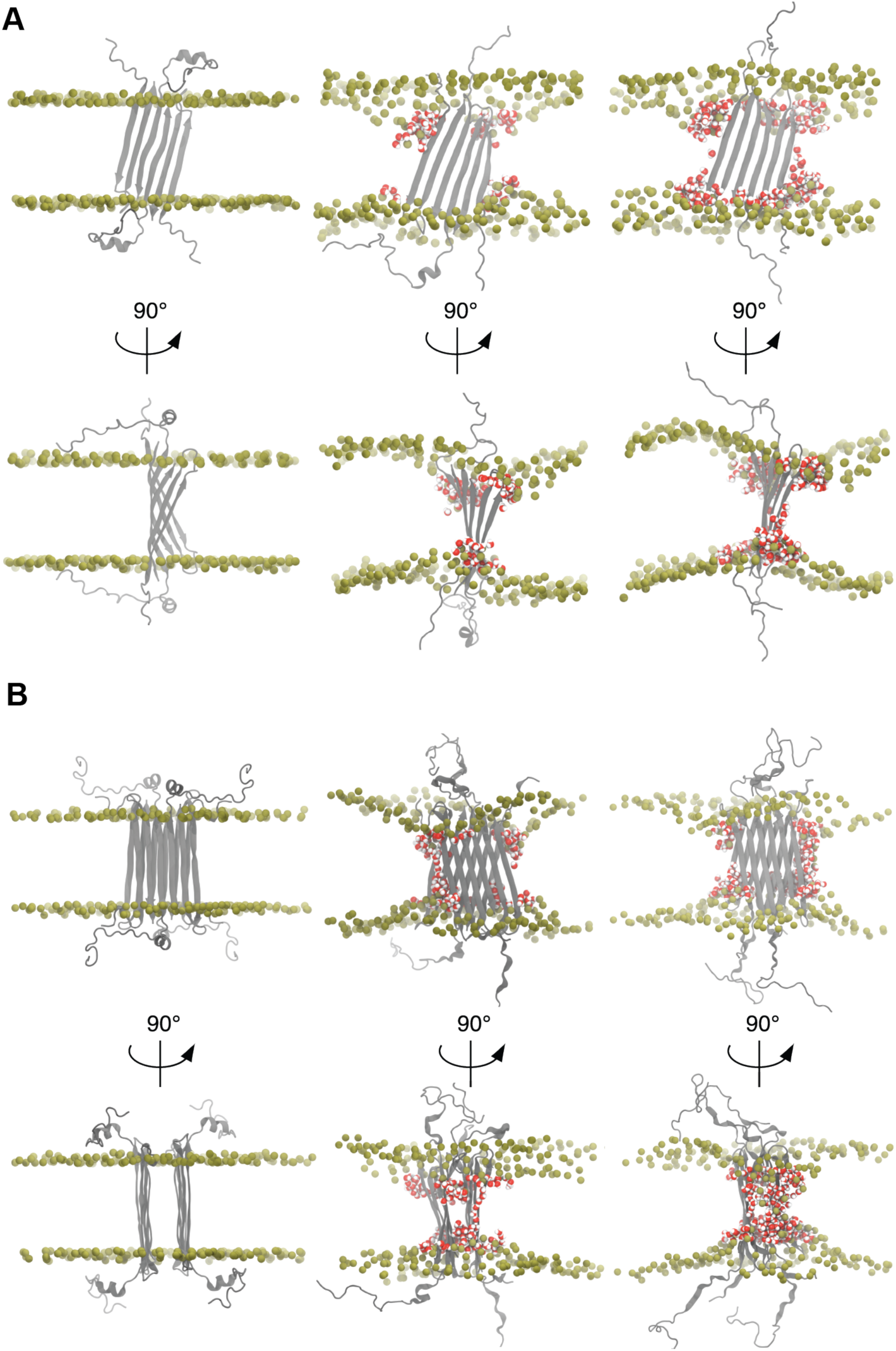
MD simulations in DPPC membrane bilayer of Aβ(1-42) (**A**) tetramer and (**B**) octamer. The snapshots shown correspond to the initial coordinates (left), after 100 ns NPT equilibrium simulation (middle), and after 100 ns NVT simulation with 100 mV applied electric field (right). Protein is shown in grey, DPPC headgroup phosphorous atoms are shown in tan, and water in red/white.

## Discussion and Conclusion

To date, only the 3D structures of Aβ fibrils had been described (*4*-*7*) and no experimental structure had been reported for Aβ oligomers, only models (*19*-*24*). Therefore, our work widens the description of the much-needed low energy structural landscape of Aβ from Aβ fibrils. This landscape evolves from the intermolecular formation of parallel β-sheets in Aβ fibrils, to intramolecular and intermolecular antiparallel β-sheet formation in the membrane-associated Aβ(1-42) oligomers reported in this work. By establishing the structure of membrane-associated Aβ(1-42) tetramers and octamers and assessing their activity in planar lipid bilayers and through MD simulations, we have revealed that their toxicity arises from the hydrophilic residues located on the edges of the β-sheets, which lead to the formation of lipid-stabilized pores. Such behavior resembles the toroidal pore-type behavior shown by many antimicrobial peptides (*25*) and would be consistent with the reported antimicrobial activity for Aβ (*26, 27*). In summary, the present work represents the resolution of the first atomic structure of an Aβ membrane-associated oligomer and describes formation of lipid-stabilized pores as the potential mechanism underlying Aβ toxicity and its relation with AD.

## Supporting information

materials and methods, supl figures and references

## References

1. D. J. Selkoe, J. Hardy EMBO Mol. Med. 8, 595–608 (2016).

2. C. Haass C. Kaether G. Thinakaran S. Sisodia Cold Spring Harb. Perspect. Med. 2, a006270–a006270 (2012).

3. I. Benilova E. Karran B. De Strooper, Nat. Neurosci. 15, 349–357 (2012).

4. X.-C. Bai et al., Nature. 525, 212–217 (2015).

5. M. A. Wälti et al., Proc. Natl. Acad. Sci. USA. 113, E4976–E4984 (2016).

6. M. T. Colvin et al., J. Am. Chem. Soc. 138, 9663–9674 (2016).

7. L. Gremer et al., Science. 358, 116–119 (2017).

8. N. Arispe E. Rojas H. B. Pollard, Proc. Natl. Acad. Sci. USA. 90, 567–571 (1993).

9. Y. Hirakura M. C. Lin, B. L. Kagan, J. Neurosci. Res. 57, 458–466 (1999).

10. H. Lin R. Bhatia R. Lal The FASEB Journal. 15, 2433–2444 (2001).

11. S. M. Butterfield, H. A. Lashuel, Angew. Chem. Int. Ed. 49, 5628–5654 (2010).

12. B. R. Roberts et al., Brain. 140, 1486–1498 (2017).

13. M. Serra-Batiste et al., Proc. Natl. Acad. Sci. U.S.A. 113, 10866–10871 (2016).

14. T. Cierpicki J. Otlewski J. Biomol. NMR. 21, 249–261 (2001).

15. A. Laganowsky E. Reading J. T. S. Hopper, C. V. Robinson, Nat. Protoc. 8, 639–651 (2013).

16. A. Leitner et al., Proc. Natl. Acad. Sci. USA. 111, 9455–9460 (2014).

17. E. Sitkiewicz J. Oledzki J. Poznanski M. Dadlez PLoS ONE. 9, e100200–14 (2014).

18. J. Marcoux et al., Structure. 22, 781–790 (2014).

19. L. Yu et al., Biochemistry. 44, 15834–15841 (2005).

20. S. Chimon et al., Nat. Struct. Mol. Biol. 14, 1157–1164 (2007).

21. M. Ahmed et al., Nat. Struct. Mol. Biol. 17, 561–567 (2010).

22. A. Laganowsky et al., Science. 335, 1228–1231 (2012).

23. D. Huang et al., J. Mol. Biol. 427, 2319–2328 (2015).

24. S. Parthasarathy et al., J. Am. Chem. Soc. 137, 6480–6483 (2015).

25. Z. O. Shenkarev et al., Biochemistry. 50, 6255–6265 (2011).

26. S. J. Soscia et al., PLoS ONE. 5, e9505 (2010).

27. D. K. V. Kumar et al., Science Translational Medicine. 8, 340ra72–340ra72 (2016).

