## Supplementary material for "Aβ(1-42) tetramer and octamer structures reveal edge pores as a mechanism for membrane damage": materials and methods, supl figures and references

*Reagents*

Lipids and detergents were purchased from Avanti Polar Lipids or Affymetrix-Anatrace. Deuterated reagents were purchased from Cortecnet or Eurisotop. All other reagents were supplied by Sigma-Aldrich unless otherwise stated. Kits for selective isotopically labeled samples were purchased from NMR-Bio. All buffers and solutions were freshly prepared using water provided by a Milli-Q system (18 MW/cm at 25ºC, Millipore).

*Purification of synthetic Aβ samples and Aβ mixtures*

Aβ(1-42) and Aβ(17-42) were synthesized and purified by Dr. James I. Elliott (New Haven, CT, USA). The following protocol, described for Aβ(1-42) but applicable to the other Aβ peptides under study, was used to obtain Aβ in a monomeric state. 5 to 10 mg of lyophilized synthetic Aβ(1-42) peptide were resuspended in 6.8 M guanidinium thiocyanate (GdnSCN) to a final concentration of 2.5 mg Aβ(1-42)/mL and sonicated for 5 min in an ice bath. Afterwards, the sample was further diluted with Milli-Q water to 1.5 mg Aβ(1-42)/mL and 4 M GdnSCN, and centrifuged. Finally, 2 mL of the 1.5 mg Aβ(1-42)/mL solution was injected into a HiLoad Superdex 30 prep grade column (GE Healthcare), previously equilibrated with 50 mM ammonium carbonate. The fractions corresponding to monomeric Aβ(1-42) were collected and their purity and concentration were determined by Reversed Phase High Performance Liquid Chromatography (RP-HPLC). The pool was finally aliquoted in the desired amounts, freeze-dried, and kept at -20ºC until use.

For mixed samples containing different peptides such as Aβ(1-42)/Aβ(17-42), we purified, as described above, the most insoluble peptide first and prepared aliquots in the desired amounts, froze them with liquid nitrogen, and kept them at -20ºC. Afterwards the second peptide was purified in the same way, aliquots were added on top of the already frozen one and the combined aliquot was freeze-dried. Samples were kept at -20ºC until use.

*Expression and purification of recombinant Aβ samples*

For all labeling schemes, Rosetta (DE3) pLysS *E. coli* cells (Novagen) were transformed with the expression vector and grown overnight at 37°C on Luria Bertani (LB)-agar plates containing 1% glucose. All cell cultures were also supplemented with 35 μg/mL chloramphenicol and 50 μg/mL kanamycin. To enhance protein production, all Aβ peptides were expressed with the construct $(His)_6$-SUMO-Aβ using SUMO as a fusion partner. An auto-induction procedure was used to produce [U-$^{15}$N] Aβ(1-42) and [U-$^{15}$N] Aβ(17-42) , as previously described (*1*). Briefly, single colonies were picked and grown overnight in LB supplemented with 1% glucose. The pre-culture was centrifuged, and the pellet was transferred to $^{15}$N-labeled P-5052 auto-inducing media with the appropriate antibiotics. The resulting cultures were grown for 6 h at 37ºC. The temperature was then lowered to 25ºC, and the culture was incubated for a further 22 h. The cells were then harvested by centrifugation and frozen at -80°C.

M9 minimal medium was used to produce [U-$^{2}$H,$^{13}$C,$^{15}$N] Aβ(1-42) and [U-$^{2}$H,$^{15}$N] Aβ(1-42) following previously reported protocols (*1*). Briefly, single colonies were picked and grown overnight in LB supplemented with 1% glucose. Cells containing the DNA construct were adapted to grow in minimal medium in a stepwise manner by inoculating the cells into fresh M9 minimal medium containing increasing percentages of $D_2O$. The final pellet, already grown overnight in M9 minimal medium prepared using 100% $D_2O$, was re-suspended and inoculated in 1 L M9 medium also prepared using 100% $D_2O$ and containing 1 g/L $^{15}NH_4Cl$ and 2 g/L D-glucose-$^{13}C_6$-1,2,3,4,5,6,6-$d_7$ or D-glucose-1,2,3,4,5,6,6-$d_7$. The culture was grown at 37°C and induced at an $OD_{600}$ ~1 by the addition of IPTG to a final concentration of 0.5 mM. After overnight growth at 25°C, the cells were harvested by centrifugation and then frozen at -80°C.

For the production of selectively labeled Ile-[$^{13}CH^3$]$^{\delta 1}$, Ala-[$^{13}CH_3$], Leu/Val-[$^{13}CH_3$]$^{proR}$ Aβ(1-42) samples, we followed previously published procedures (*2*). Briefly, 2-[$^{13}CH_3$], 4-[$^{2}H_3$] acetolactate (NMR-Bio) at 300 mg/mL was added 1 h prior to induction. Forty minutes later (20 min prior to induction), 2-hydroxy-2-(1'-[$^{2}H_2$], 2'-[$^{13}$C])ethyl-3-keto-4-[$^{2}H_3$]butanoic acid (NMR-Bio) at 60 mg/mL and 2-[$^{2}$H], 3-[$^{13}$C]alanine (NMR-Bio) at 700 mg/mL were added. Protein expression was induced with IPTG as previously described.

The following protocol, described for Aβ(1-42) but applicable to the other Aβ peptides under study, was used to purify and obtain Aβ in a monomeric state. After protein expression, cells were lysed by sonication and centrifuged, and the supernatant was then purified as already described (*1*). Briefly, the cleared soluble fraction was loaded onto a HisTrap HP 5-mL Ni column (GE Healthcare) and the fusion protein was eluted with 0.5 mM imidazole. IMAC fractions were analyzed by sodium dodecyl sulfate polyacrylamide gel electrophoresis (SDS-PAGE), and those containing the fusion protein were pooled. Next, the buffer was exchanged using a HiPrep 26/10 desalting column (GE Healthcare) equilibrated with 50 mM ammonium carbonate and 1 mM TCEP. Afterwards, the concentration and purity of protein was determined by Nanodrop® and RP-HPLC. Subsequently, samples were incubated overnight at 4ºC with SUMO protease (Ulp1) in a 1:50 protease:protein ratio to cleave Aβ(1-42) from the SUMO fusion tag. The concentration of Aβ(1-42) peptide after the cleavage was determined by RP-HPLC analysis. Subsequently, aliquots containing 3.75 mg Aβ(1-42) were prepared and freeze-dried. Each of these aliquots was solubilized with 6.8 M GdnSCN to 2.5 mg Aβ(1-42)/mL and sonicated for 5 min in an ice bath. Afterwards, the sample was further diluted with Milli-Q water to 1.5 mg Aβ(1-42)/mL and 4 M GdnSCN, and then centrifuged. Finally, 2.5 mL of the 1.5 mg Aβ(1-42)/mL solution was injected into a HiLoad Superdex 30 prep grade column (GE Healthcare), previously equilibrated with 50 mM ammonium carbonate. The peaks corresponding to SUMO and monomeric Aβ(1-42) were collected separately, and their purity and concentration were determined by RP-HPLC. The pool containing pure Aβ(1-42) was aliquoted in the desired amounts, freeze-dried, and kept at -20ºC until use.

*Isotope-labeled samples*

The following samples were produced: [U-$^{15}$N]-Aβ(1-42), [U-$^{15}$N]-Aβ(17-42) for 2D $^{1}$H-$^{15}$N-HSQC experiments; [U-$^{2}$H,$^{13}$C,$^{15}$N]-Aβ(1-42) for 3D backbone assignment experiments; [U-$^{2}$H,$^{13}$C,$^{15}$N]-Ile-[$^{13}$CH$^{3}$]$^{\delta1}$, Ala-[$^{13}CH_3$], Leu/Val-[$^{13}CH_3$]$^{proR}$-Aβ(1-42) for side chain methyl assignments; and [U-$^{2}$H,$^{15}$N]-Ile-[$^{13}$CH$^{3}$] $^{\delta1}$, Ala-[$^{13}CH_3$], Leu/Val-[$^{13}CH_3$]$^{proR}$-Aβ(1-42) for 3D $^{13}CH_3$-$^{13}CH_3$ and NH-$^{13}CH_3$ NOESY experiments.

*Preparation of βPFOs$_{Aβ(1-42)}$ sample for NMR experiments*

For NMR experiments, *βPFOs$_{Aβ(1-42)}$* were prepared by dissolving lyophilized isotopically labeled monomeric Aβ(1-42) in the required volume of 10 mM Tris-$d_{11}$ and 28.5 mM DPC-$d_{38}$ to reach 1 mM Aβ(1-42). Afterwards, the pH was checked and adjusted to pH 9.5 or 8.5 with either a 10% HCl or a 10% NaOH solution, and the sample was left incubating at 37°C for 24 h. To prepare *βPFOs$_{Aβ42}$* at other Aβ(1-42) concentrations, only the concentration of DPC micelles ([$M_{DPC}$]) was adjusted so that the final [Aβ(1-42]/[$M_{DPC}$] ratio was 2:1, where [$M_{DPC}$] is the concentration of DPC micelles and equals to the difference between the DPC detergent concentration ([$D_{DPC}$]) and its critical micellar concentration (CMC) divided by its aggregation number (*i.e.* ([$D_{DPC}$]-CMC)/aggregation number)). The CMC of DPC was taken to be 1.5 mM (*3*, *4*) and the DPC aggregation number 54 (*4*).This preparation, which is later referred to in the paper as *βPFOs$_{LOW_A\beta(1-42)}$,* was found to be enriched mainly in Aβ(1-42) tetramers.

<u>*NMR experiments*</u>

All experiments were carried out at 37°C on a 900 MHz Bruker Avance III HD spectrometer equipped with a 5-mm CP-TCI cryogenic probe, or an 800 MHz Bruker Avance III HD spectrometer equipped with a 3-mm CP-TCI cryogenic probe, both instruments located at the Swedish NMR Centre in Gothenburg, or an 800 MHz Bruker Avance III HD spectrometer equipped with a 5-mm CP-TCI cryogenic probe, located at the IECB in Bordeaux.

For the resonance assignment of backbone atoms of the Aβ(1-42) tetramer prepared in DPC micelles, experiments from the standard Bruker library were recorded (HNCA, HNCACB, HNCO and HN-NH NOESY). For the resonance assignment of methyl groups, experiments from the standard Bruker library were recorded (Hme)Cme([C]CA)CO, (Hme)Cme([C]CA)NH, Hme(Cme[C]CA)NH (*5*) and complemented with (H)C-TOCSY-C-TOCSY-(C)H experiments (*6*). Additionally, four 3D SOFAST-NOESY-HMQC experiments (*5*) were recorded to obtain NOE correlation between methyl groups (Hm-HmCm and Cm-HmCm) and between methyl and amide protons (Hm-NH and Cm-HN) of the Aβ(1-42) tetramer in DPC micelles. The acquisition parameters for all the NMR experiments carried out are summarized in Table S3. All these experiments were acquired using non-uniform sampling (NUS) and processed with *mddnmr* software (*6*, *7*).

<u>*NMR amide temperature coefficients*</u>

Amide temperature coefficients of the Aβ(1-42) tetramer were determined by measuring the 2D [$^{1}$H,$^{15}$N]-TROSY spectra of the Aβ(1-42) tetramer sample at 303 K, 310 K, 317 K, and 324 K on a 900 MHz Bruker Avance III HD spectrometer equipped with a 5-mm CP-TCI cryogenic probe, and calculated using the following equation:

$$Temperature\ coefficient\ values = {}^{\Delta\delta_{NH}}/_{\Delta T}$$

It is well established that, in aqueous solvents, exposed NHs typically display gradients from -6 to -8.5 ppb/K (*8*) while hydrogen-bonded exchange-protected NHs are characterized less negative Δδ/ΔT values than -4 ppb/K. However, numerous exceptions to these generalizations occur. In our case, the first anomalous observation was the fact that all amide protons presented positive Δδ/ΔT, meaning that chemical shifts of amide proton resonances shift downfield as the temperature increases. These downfield shifts with increased temperature may be explained by greater solvent protection of NH protons (*8*), probably due to the effect of the detergent micelle surrounding the tetramer. Furthermore, we observed that most of the NH amide protons of residues from β1, β2 and β3 were the most affected by temperature changes. Cierpicki *et al.* noted that amides involved in hydrogen bonds with a length of less than approximately 3.0 Å exhibited a larger temperature coefficient, because the secondary chemical shift caused by hydrogen bonding is greater, and so the same fractional change gives rise to a larger gradient (*9*). This report would be consistent with residues with the largest $\Delta\delta/\Delta T$ being involved in stable hydrogen bonds.

*NMR titrations with paramagnetic reagents*

NMR titrations with 16-DOXYL-stearic acid (16-DSA) were performed by addition of concentrated stock solutions of 16-DSA to an Aβ(1-42) tetramer NMR sample. Stock solutions of 16-DSA were obtained by dissolving this chemical in methanol-$d_4$. The Aβ(1-42) tetramer NMR sample was prepared at 1 mM Aβ(1-42), using the appropriately labeled sample, in 10 mM Tris-$d_{11}$, 28.5 mM DPC-$d_{38}$ at pH 9.5. 16-DSA stock solution was subsequently added to this Aβ(1-42) sample to obtain the following concentrations of 16-DSA: 0.3, 0.45, 0.6, 1.2 and 2.4 mM. [$^1$H,$^{15}$N]-TROSY and [$^1$H,$^{13}$C]-HMQC experiments were acquired at each 16-DSA concentration point. These NMR experiments were also performed on a sample without 16-DSA in order to be used as reference.

*NMR structure calculation of the Aβ(1-42) tetramer*

The structure of the Aβ(1-42) tetramer was determined with the iterative ARIA2.3/CNS1.21 software (*11*, *12*). Distance restraints were derived from NOE cross-peaks (HN-HN, HN-Methyl, Methyl-Methyl) and used as input for ARIA, together with dihedral angle restraints and hydrogen-bond restraints. Upper bound distances for NOE restraints were derived from NOE cross-peaks volumes using characteristic distances (sequential NH-NH NOEs in β-sheet or intra-residual NH-Methyl in alanines). Backbone dihedral angles were predicted from backbone chemical shifts with TALOS-N (*13*). Predictions classified as "Strong" were converted to dihedral angle restraints with an error corresponding to twice the standard deviation given by TALOS-N. Hydrogen bond restraints for anti-parallel beta-strand pairing were deduced from the NOE pattern and confirmed by initial calculations from NOEs and dihedral angle restraints only. Each hydrogen bond is encoded by two restraints (HN…O with upper-bound 2.3 Å and N…O, upper-bound 3.3 Å). During structure calculation, four copies of an Aβ(1-42) chain were modeled, using NCS restraints to maintain each dimer superimposable in the tetramer. For each iteration, 100 conformations were generated, except for the last iteration, where 500 conformers were calculated. The 50 lowest-energy conformers were refined in a shell of DMSO molecules (*14*) and the 15 refined conformers with the least number of distance restraint violations were selected as the final Aβ(1-42) tetramer ensemble. The structure ensemble was validated with PROCHECK (*15*) WHATIF (*16*) and MolProbity (*17*).

*Simulations of Aβ(1-42) tetramer solubilization in DPC*

From the NMR ensemble of Aβ(1-42) tetramer structures determined with ARIA, the conformer with the least restraint violations was selected to be solubilized in n-dodecylphosphocholine (DPC) micelle. The SimShape protocol (*18*) was used to accelerate the assembly of the protein-micelle complex. This method uses grid-steered molecular dynamics to assemble detergents into a toroidal micelle that wraps around the hydrophobic core of a membrane protein. Three biasing potentials were utilized in total. Two took the shape of concentric toroids, where the first toroid was coupled to the head-group heavy atoms of DPC, while the second smaller torus was coupled to the tail-heavy atoms. The third biasing potential took the shape of a plane and was coupled to the detergent tail-heavy atoms.

Prior knowledge of the approximate micelle shape and orientation facilitated the design of the shape of the toroidal potentials. HDX studies characterized the central six-stranded β-sheet region of the tetramer with slow water exchange, suggesting burial of these residues within the micelle. This ~29 Å long region of the Aβ(1-42) tetramer was used to inform the dimensions of the toroid-shaped grid potentials. The number of DPC molecules in the micelle was determined to be 120 by estimating the size of the tetramer-micelle complex using overall correlation time obtained from NMR (*10*). The DPC molecule positions were initialized in a toroidal pattern around the protein such that there was a minimum distance of 10 Å between any given pair of detergent and protein atoms.

The complex was simulated using NAMD with grid-steered molecular dynamics for 1 ns at 310 K with a Langevin thermostat with a damping coefficient of 5/ps. Generalized Born Implicit Solvent (GBIS) with ionic strength 0.15 mM and dielectric constant of 80 were used. CHARMM36 forcefield parameters were used. Head and tail atoms were separately coupled to their grid potentials with scaling factors of 0.18 and 0.25, respectively. The tail atoms were additionally coupled to the planar grid potential with a scaling factor of 0.14. Backbone heavy atoms were harmonically restrained using a 1 kcal/(mol·Å$^2$) spring constant to maintain protein structure during detergent assembly. After 1 ns, detergent molecules assembled into a micelle around the protein with a toroidal shape similar to the attractive grid potentials (fig. S13).

Continuing with the SimShape protocol, the micellar complex was completely uncoupled from the toroidal attractive potentials, solvated with TIP3P water, and ionized with 150 mM NaCl. The solvated system was minimized with conjugate gradient energy minimization for 5,000 steps, and simulated for 30 ns with protein backbone heavy atom harmonic position restraints with a spring constant of 1 kcal/(mol·Å$^2$). Next, the protein restraints were gradually removed to allow for complete equilibration of the complex (Fig. 2E). The spring constant was lowered in steps by 0.001 kcal/(mol·Å$^2$) every 10 ps for 10 ns. Finally, eight replicates of the unrestrained protein-micelle complex were simulated for an additional 100 ns each with no biasing forces applied. These equilibrium trajectories were then used to characterize contacts between the tetramer and detergents.

Contacts were calculated between the Aβ(1-42) tetramer and DPC molecules and are shown in fig. S14. Two contact sites on DPC were considered: the head group nitrogen atom and the terminal tail carbon atom. The number of contacts was defined as the number of contact sites within 9 Å of an amide backbone nitrogen atom and was calculated for every frame, summed over symmetric chains, and averaged over a 100-ns trajectory, returning an average number of contacts per residue. The per-residue average contact number of the eight independent replicates was then considered to be eight independent samples, allowing for calculation of statistical error per residue, shown as +/- standard deviation divided by the number of independent samples.

*Preparation of $\beta PFOs_{LOW\_A\beta(1-42)}$ and $\beta PFOs_{HIGH\_A\beta(1-42)}$ samples*

$\beta PFOs_{LOW\_A\beta(1-42)}$ and $\beta PFOs_{HIGH\_A\beta(1-42)}$ corresponding to Aβ(1-42]/[$M_{DPC}$] ratios of 2:1 and 6:1, respectively, were prepared from freeze-dried monomeric Aβ(1-42) samples and dissolved in 10 mM Tris, 5.5 mM DPC adjusted to pH 9, reaching a final concentration of 150 µM Aβ(1-42) in the case of $\beta PFOs_{LOW\_A\beta(1-42)}$ and 450 µM Aβ(1-42) in the case of $\beta PFOs_{HIGH\_A\beta(1-42)}$. The samples were incubated at 37ºC for 24 h.

*SEC*

*βPFOs*$_{LOW\_A\beta(1\text{-}42)}$ and *βPFOs*$_{HIGH\_A\beta(1\text{-}42)}$ samples were injected into a tandem Superdex 200 increase 10/300 (GE Healthcare). The columns were equilibrated with 10 mM Tris, 100 mM NaCl at pH 9 containing 3 mM DPC and eluted at 4ºC at a flow rate of 0.5 mL/min.

<u>SEC/IM-MS</u>

An Acquity UPLC H-class system (Waters, Manchester, UK) comprising a quaternary solvent manager, a sample manager set at 10°C, a column oven and a TUV detector operating at 280 nm and 214 nm was coupled to a Synapt G2 HDMS mass spectrometer (Waters) for online SEC/IMS-MS instrumentation (*20*). An Acquity BEH SEC column (4.6x150 mm, 1.7 µm particle size, 200 Å pore size) (Waters) was equilibrated with 200 mM $(NH_4)_2CO_3$, 14.2 mM $C_8E_5$ at pH 9.0 and run with the following flow rate gradient: 0.25 mL/min over 4 min; then 0.10 mL/min over 6 min and finally 0.25 mL/min over 2 min.

The IMS-MS experiments are reported as recommended by Gabelica *et. al.* (*21*). The Synapt G2 HDMS was operated in positive mode with a capillary voltage of 3.0 kV. The main parameters—sample cone 180 V, trap collision energy 100 V, trap gas flow 5 mL/min and backing pressure 6 mbar—were finely tuned to disrupt the detergent protein interaction and to maintain the oligomer species. Acquisitions were performed in the m/z range 1000–10,000 with a 1.5-s scan time. External calibration was performed using singly charged ions produced by a 2 g/L solution of cesium iodide dissolved in 2-propanol/water (50/50, v/v). To assess the effect of activation energy on the oligomeric species detected, we increased the energy conditions by means of the sampling cone (100 V, 180 V and 200 V) and the trap collision energy (50 V, 100 V, and 160 V). MS data interpretations were performed using Mass Lynx V4.1 (Waters).

In order to unambiguously assign the oligomeric state of each MS peak and to measure the drift time of each species, an ion mobility method using travelling wave IMS (TWIMS) technology was optimized as described below. Prior to TWIMS separation, ions were thermalized in the helium cell (180 mL/min). Subsequently, ion separation was performed in the pressurized ion mobility cell using a constant $N_2$ (purity >99%) flow rate of 90 mL/min. The wave height and velocity were 40 V and 800 m/s, respectively. Transfer collision energy was set to 15 V to extract the ions from the IM cell to the TOF analyzer. IMS-MS experiments were performed in triplicate under identical instrumental conditions. We assessed the CCS from the mobility measurements using Equation 1 (*22*):

$$\mathrm{CCS} = \frac{3}{16}\sqrt{\frac{2\pi}{\mu k_B T}}\frac{ze}{NK} = \frac{3}{16}\sqrt{\frac{2\pi}{\mu k_B T}}\frac{ze}{N_0\left(\frac{p}{p_0}\frac{T_0}{T}\right)K} = \frac{3}{16}\sqrt{\frac{2\pi}{\mu k_B T}}\frac{ze}{N_0 K_0} \quad \text{(Equation 1)}$$

IMS data were calibrated to perform CCS calculations using the most intense charge state of external calibrants prepared under non-denaturing conditions. The choice of calibrants is a critical point in the calibration framework (*11*) and even more when working with membrane proteins because of the reduced charge states and lower mobility (*12*). Allison *et. al.* put emphasis on the relation of CCS/*z* (CCS'), which is substantially increased in the case of membrane proteins as a result of their reduced charge. For this reason, they propose the use of large soluble proteins as calibrants to obtain CCS' closer to the values of the membrane protein under study. In Table S4, we show the reported CCS' values for the calibrants used in this study (*11*) and the theoretical CCS' values calculated for the tetramer structure and the derived models. Theoretical CCS' for our samples were derived from PDB files corresponding to the NMR structure of the Aβ(1-42) tetramer and to two octamer models built using the structure of the Aβ(1-42) tetramer as a building block. The first octamer model was based on the association of two tetramers to form a loose β-barrel structure and the second one on the association of two tetramers in a β-sandwich structure. Theoretical CCS value for the Aβ(1-42) tetramer and octamers were obtained using the Projection Approximation method within the Impact software (*25*). Their corresponding PDB files were used as an input file and were run on Impact with a convergence value of 1% enabled to determine an average CCS as a mean of three independent calculations. Solvent heteroatoms were excluded from the theoretical calculation.

*Cross-linking of* $\beta PFOs_{LOW\_A\beta(1\text{-}42)}$ and $\beta PFOs_{HIGH\_A\beta(1\text{-}42)}$ *samples*

$\beta PFOs_{LOW\_A\beta(1\text{-}42)}$ and $\beta PFOs_{HIGH\_A\beta(1\text{-}42)}$ were prepared as described in the section "Preparation of $\beta PFOs_{LOW\_A\beta(1\text{-}42)}$ and $\beta PFOs_{HIGH\_A\beta(1\text{-}42)}$", with the exception that 10 mM sodium carbonate ($Na_2CO_3$) was used instead of 10 mM Tris to avoid the interference of Tris in the cross-linking reaction. After sample preparation, the concentration of $\beta PFOs_{LOW\_A\beta(1\text{-}42)}$ was maintained at 150 μM Aβ(1-42) while that of $\beta PFOs_{HIGH\_A\beta(1\text{-}42)}$ was brought from 450 μM to 150 μM Aβ(1-42) by diluting it with a solution containing 10 mM $Na_2CO_3$, 1.5 mM DPC at pH 9. Afterwards, both samples were cross-linked using (4-(4,6-dimethoxy-1,3,5-triazin-2-yl)-4-methyl-morpholinium chloride) (DMTMM) as the cross-linking reagent (*13*). DMTMM was added to a final concentration of 15 mM (4.15 mg/mL) and the samples were incubated for 2 h at 50°C and 800 rpm. Samples were quenched by directly preparing them for SDS-PAGE and high-mass MALDI analysis.

*SDS-PAGE analysis of cross-linked βPFOs$_{LOW_Aβ(1-42)}$ and βPFOs$_{HIGH_Aβ(1-42)}$ samples*

The cross-linked samples were diluted to 50 µM Aβ(1-42) using a solution of 10 mM $Na_2CO_3$ and 1.5 mM DPC at pH 9. Finally, 20 µL of the resulting solution was mixed with 10 µL of 3X sample buffer (3X SB) and 20 µL of the mixture, either non-boiled or boiled (for 5 min at 95°C) were electrophoresed in 1 mm-thick SDS-PAGE gels containing 15% acrylamide. Gels were run at 50 V for 30 min, 120 V for 2 h and stained using Coomassie Blue.

*High-mass MALDI-MS analysis of cross-linked βPFOs$_{LOW_Aβ(1-42)}$ and βPFOs$_{HIGH_Aβ(1-42)}$ samples*

Before MALDI-TOF analysis, cross-linked samples were diluted down to 37.5 µM Aβ(1-42) in $H_2O$. This dilution step is critical to reduce the amount of detergent that could later interfere with the co-crystallization of the sample with the matrix, which is an essential prerequisite for the MALDI ionization process (*14*). Next, diluted samples were mixed (1:1 v/v) with a matrix solution of sinapic acid (10 mg/ml) containing (1:1 v/v) acetonitrile/deionized water with 0.1% trifluoroacetic acid (TFA). Each mixture (2 µl thereof) was deposited on the MALDI target plate using the dried-droplet method. As a control, 2 µl of *βPFOs$_{LOW_Aβ(1-42)}$* and *βPFOs$_{HIGH_Aβ(1-42)}$* samples prepared as described but without adding the cross-linking reagent were examined using the same deposition method. High-mass MALDI-MS analyses were carried out on a MALDI-TOF mass spectrometer (Autoflex III, Bruker) used in linear mode and equipped with a HM3 high-mass detector (CovalX AG), which allows the (sub-µM) detection of macromolecules up to 1500 kDa with low saturation. Calibration was achieved using singly and doubly charged bovine serum albumin ions ($[M + 2H]^{2+}$= 33216 Da and $[M + H]^{+}$= 66431 Da) and the gas phase dimer of this protein ($[2M + H]^{+}$= 132861 Da). The mass spectra were acquired by averaging 2000 shots (8 different positions into each spot and 250 shots per position), using the same laser fluency before and after crosslinking. The spectra were processed (including background subtraction and smoothing) using FlexAnalysis (Bruker).

*Electrical recordings with planar lipid bilayers*

Ionic currents from planar bilayers formed from diphytanoyl-*sn*-glycero-3-phosphocholine in 10 mM Tris·HCl and 150 mM NaCl at pH 7.5 and 23°C were measured by applying a 2 kHz low-pass Bassel filter with a 10 kHz sampling rate. Potentials were applied, and the current was recorded using Ag/AgCl electrodes connected to a patch-clamp amplifier (Axopatch 200B, Axon Instruments). Current recordings were analyzed using the Clampfit 10 software package (Molecular devices). Open-pore currents were measured by a Gaussian fit to all-point histogram. The center of the peak corresponds to the open-pore conductance and the width at half height to the error. Each electrophysiology chamber contained 500 µL 10 mM Tris·HCl and 150 mM NaCl at pH 7.5. Two samples were analyzed, $\beta PFOs_{LOW\_A\beta(1\text{-}42)}$, which was diluted from 1:250 to 1:100 in the chamber (*10*), and $\beta PFOs_{HIGH\_A\beta(1\text{-}42)}$, which was diluted 1:100 in the chamber. For $\beta PFOs_{LOW\_A\beta(1\text{-}42)}$, type 1, 2 and 3 pores were observed in 17%, 48% and 35% of the experiments (N=105). For $\beta PFOs_{HIGH\_A\beta(1\text{-}42)}$, type 1, 2, and 3 pores were observed in 8.5%, 35%, and 56% of the experiments (N=71). Controls were carried out to establish that the concentration of the detergent micelles present in the samples did not affect the stability of the bilayer.

*Simulations of the Aβ(1-42) tetramer and octamer in a DPPC bilayer*

Molecular dynamics simulations of the Aβ(1-42) tetramer and octamer in planar lipid bilayers were performed under an applied electric field using NAMD 2.13. Two structures were simulated, each in triplicate, one corresponding to the Aβ(1-42) tetramer structure obtained by NMR and the other to the octamer β-sandwich structure determined by CCS. These structures were aligned to the principal axes of their β-sheet regions, and embedded into 80Åx80Å planar 1,2-dipalmitoyl-sn-glycero-3-phosphocholine (DPPC) bilayers using the CHARMM-GUI input generator (*28*). The systems were further simulated at equilibrium NPT conditions for 100 ns with semi-isotropic pressure coupling in the X-Y plane. The systems were then equilibrated in an NVT ensemble for 10 ns. Then an external electric field of 100 mV was applied along the Z-axis and each system was simulated for another 100 ns.

Contacts between the bilayer DPPC molecules and the Aβ(1-42) tetramer and octamer during the applied 100 mV electric field were calculated. Two contact groups on DPPC were considered: the headgroup nitrogen and the two terminal tail carbon atoms. The number of contacts was defined as the number of contact sites within 9 Å of an amide backbone nitrogen atom and was calculated for every frame, summed over symmetric chains, and averaged over the 100 ns trajectory, returning an average number of contacts per residue. The per-residue average contact number of the three independent replicates was then considered three independent samples, allowing for calculation of statistical error per residue, shown as +/- standard deviation divided by the number of independent samples.

### Supporting Figures

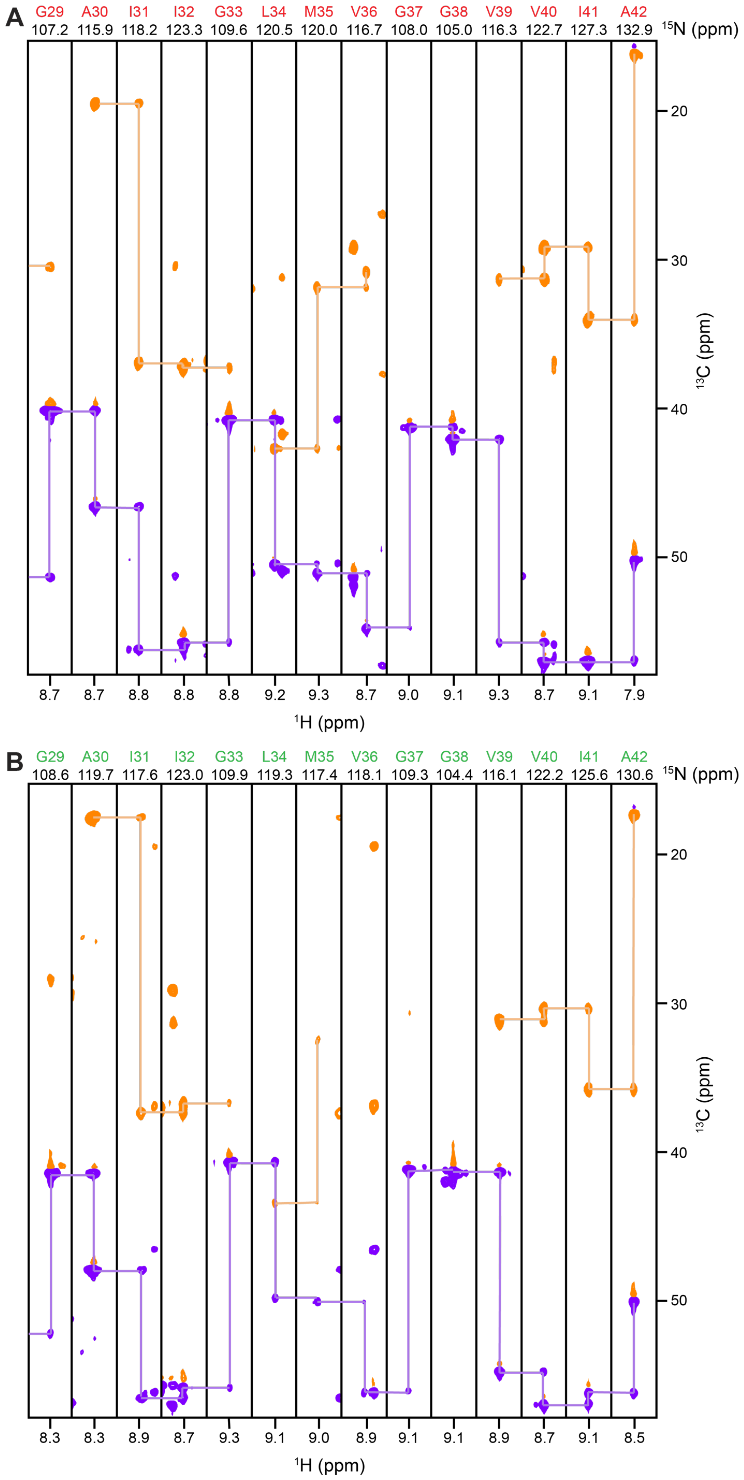


**fig. S1.** Sequential assignments for $\beta PFOs_{A\beta(1\text{-}42)}$ sample. Strips for residues 29-42 for (**A**) red Aβ(1-42) subunit and (**B**) green Aβ(1-42) subunit from a 3D TROSY-HNCACB experiment obtained using a $^{2}H,^{15}N,^{13}C$ $\beta PFOs_{A\beta(1\text{-}42)}$ sample. Purple and orange peaks arise from Cα and Cβ nuclei, respectively. The purple and orange lines indicate the sequential connection between the strips.

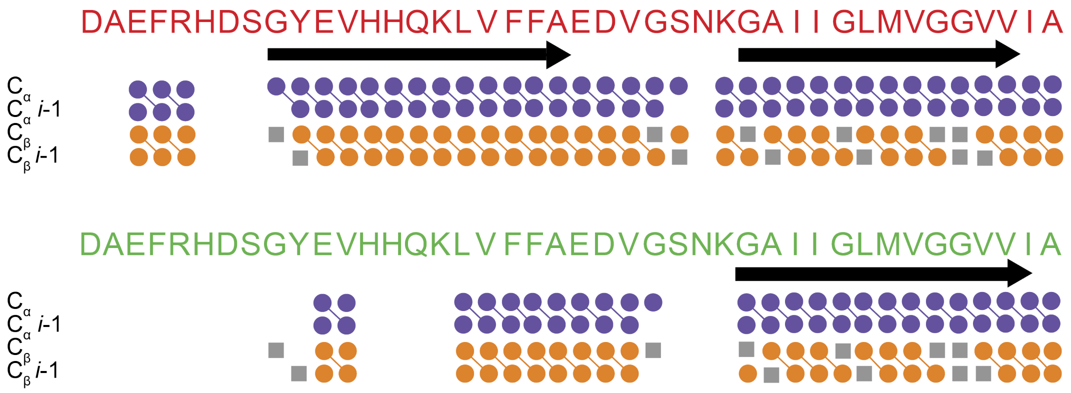


**fig. S2.** Summary of the data establishing the sequence-specific resonance assignments of *βPFOs*$_{A\beta(1\text{-}42)}$ sample. Residues for which the intraresidual Cα, sequential Cα, intraresidual Cβ, sequential Cβ were observed in the 3D TROSY-HNCA or the 3D TROSY-HNCACB are marked, respectively, as a dot in the rows "CA", "CA$_{i-1}$", "CB", "CB$_{i-1}$". Lines indicate connectivity between the atoms. Gray boxes indicate unobservable signals (Glycine Cβ). The arrows show the location of the three β-strands.

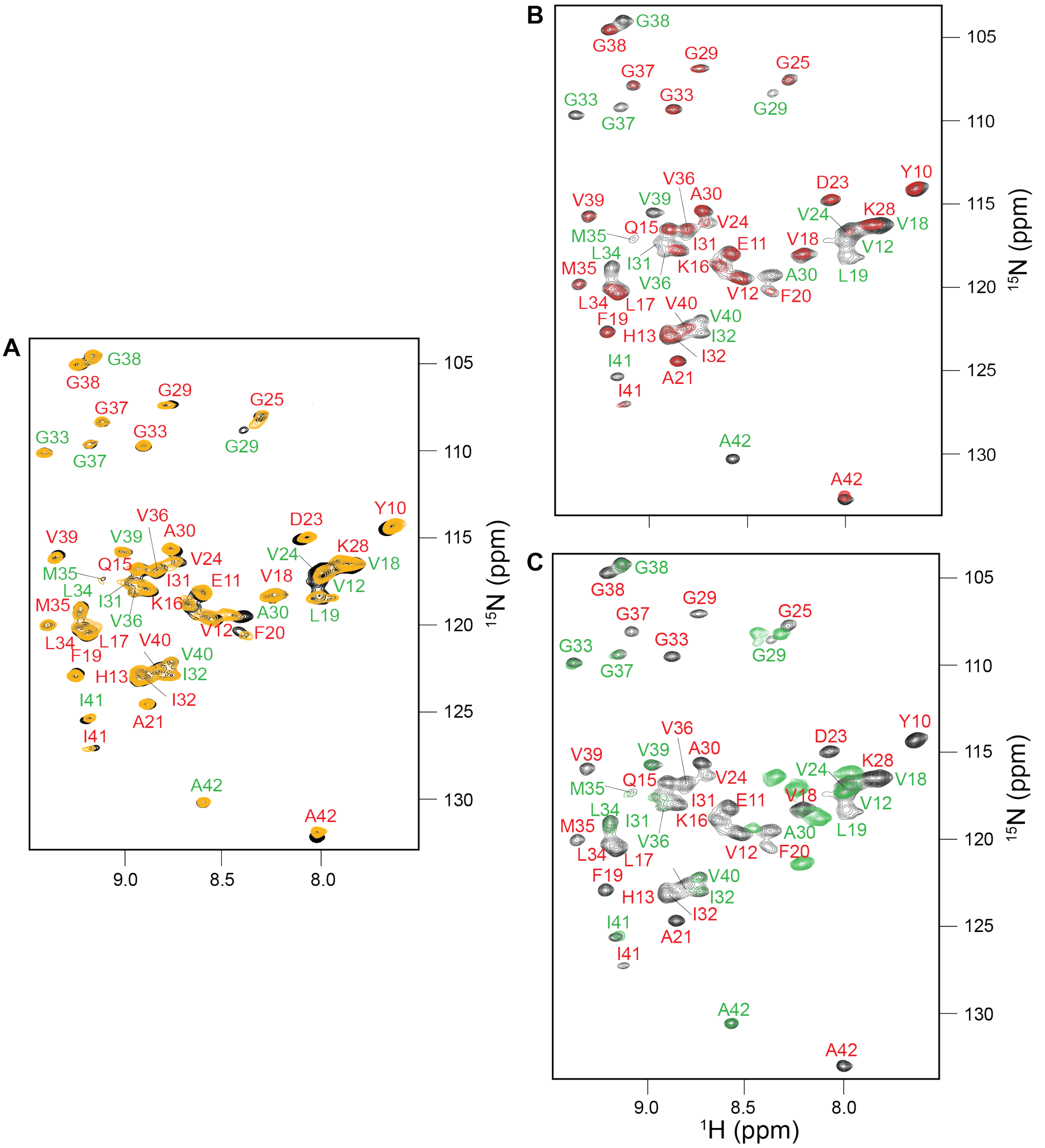


**fig. S3.** 2D [$^{1}$H,$^{15}$N]-HSQC spectra of mixtures of Aβ(1-42) and Aβ(17-42) with different labeling schemes to unambiguously establish the connectivity between β1 and β2, and α1 and β3 secondary structural elements. We hypothesized that a sample prepared using a mixture of Aβ(1-42) and Aβ(17-42) should form the same tetramer structure as Aβ(1-42) by itself, since Aβ(1-42) and Aβ(17-42) comprise, respectively, the required residues to incorporate in the tetramer as the red and green Aβ subunits. (**A**) We confirmed this hypothesis by establishing that the 2D [$^{1}$H,$^{15}$N]-HSQC spectrum of a sample prepared using a mixture of $^{15}$N Aβ(1-42) and $^{15}$N Aβ(17-42) in orange, was identical to that of a sample prepared using $^{15}$N Aβ(1-42) alone, shown in black. (**B**) Next, we measured 2D [$^{1}$H,$^{15}$N]-HSQC spectrum of a tetramer sample prepared using a mixture of $^{15}$N Aβ(1-42) and $^{14}$N Aβ(17-42) shown in red and compared it to that obtained for a sample prepared using only $^{15}$N Aβ(1-42) in black. Only residues assigned to β1 and β2 of the red Aβ subunit were detected. (**C**) Finally, we measured 2D [$^{1}$H,$^{15}$N]-HSQC spectrum of a tetramer sample prepared using a mixture of $^{14}$N Aβ(1-42) and $^{15}$N Aβ(17-42) shown in green and compared to that obtained for a sample prepared using only $^{15}$N Aβ(1-42) in black. Only residues assigned to α1 and β3 of the green Aβ subunit were observed. In summary, these results validated our assignments, providing conclusive evidence for β1 being connected to β2 within the red Aβ(1-42) subunit and α1 being connected to β3 within the green Aβ(1-42) subunit, thereby further confirming the Aβ(1-42) tetramer topology. In all 2D [$^{1}$H,$^{15}$N]-HSQC spectra, two Aβ(1-42) subunits were detected, and residues belonging to each of them were labeled in either red or green.

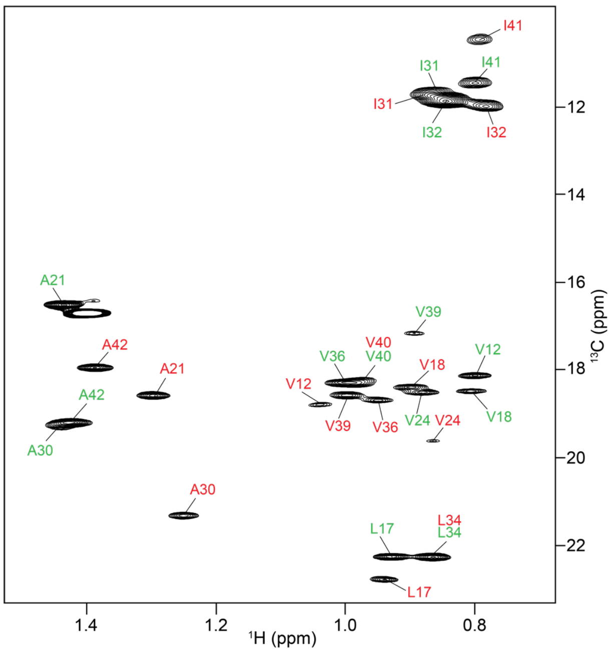


**fig. S4.** Alanine, isoleucine, leucine, and valine methyl group resonance assignments of the Aβ(1-42) tetramer. The 2D [$^{1}$H,$^{13}$C]-TROSY spectrum of the Aβ(1-42) tetramer prepared using selectively $^{13}$C methyl-protonated AILV and otherwise uniform $^{2}$H,$^{15}$N Aβ(1-42) in DPC at 37°C. Two Aβ(1-42) subunits are detected and residues belonging to each of them are labeled in either red or green. All available resonance assignments are indicated (6/8 Ala, 6/6 Ile, 4/4 Leu, and 12/12 Val).

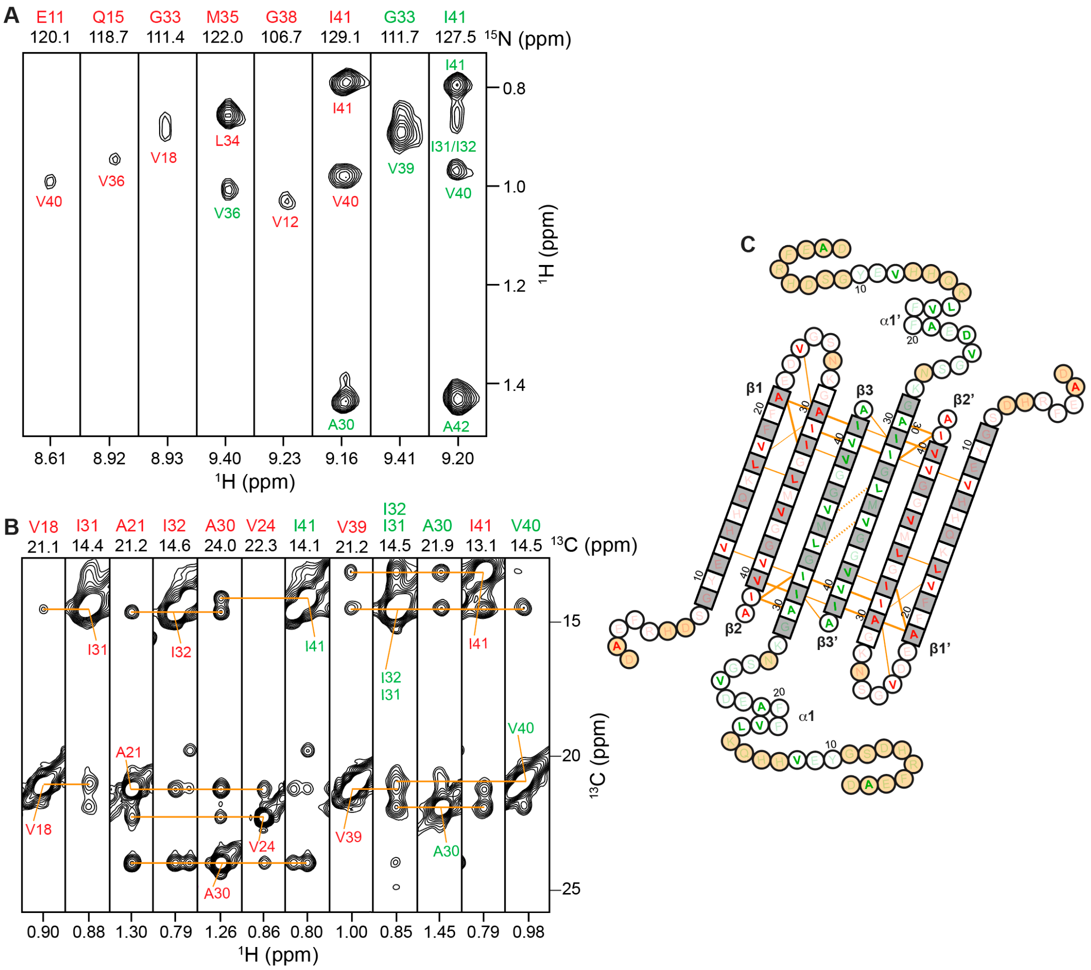


**fig. S5.** Alanine, isoleucine, leucine, and valine methyl labeling to validate Aβ(1-42) tetramer topology. (**A**) NH-CH$_3$ NOE strips from a 3D NH-CH$_3$ NOESY spectrum and (**B**) CH$_3$-CH$_3$ NOE strips from a 3D CH$_3$-CH$_3$ NOESY spectrum. Both spectra were recorded using the Aβ(1-42) tetramer prepared using selectively $^{13}$C methyl-protonated AILV and otherwise uniform $^{2}$H,$^{15}$N Aβ(1-42) in DPC at 37°C. (**C**) Aβ(1-42) tetramer topology. The color of the amino acid indicates whether it belongs to the red or green Aβ(1-42) subunit and $^{13}$C methyl-protonated AILV residues are highlighted. Amino acids in square denote β-sheet secondary structure, as identified by secondary chemical shifts; all other amino acids are in circles. Orange lines denote experimentally observed CH$_3$-CH$_3$ NOEs. For clarity, NH-CH$_3$ NOEs are not shown. Bold lines indicate strong NOEs typically observed between hydrogen-bonded residues in β-sheets. Dashed lines show probable contacts between protons with degenerate $^{1}$H chemical shifts. The side chains of white and grey residues point towards distinct sides of the β-sheet plane, respectively. Orange circles correspond to residues that could not be assigned.

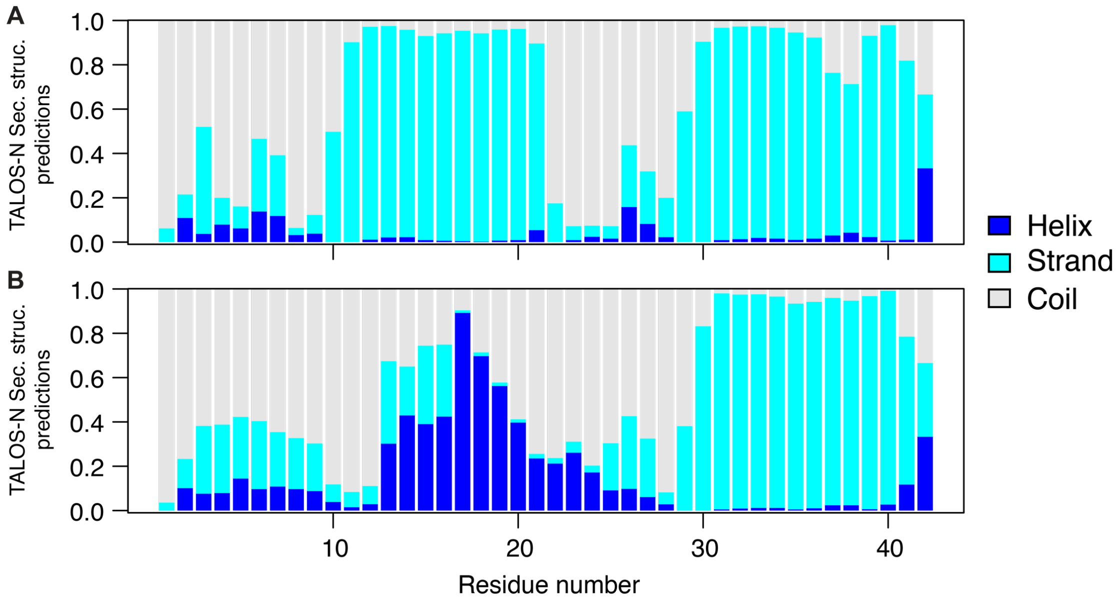


**fig. S6**. Secondary structure prediction confidence reported by TALOS+ from chemical shifts (N, Cα, C, Cβ and HN) for (**A**) the Aβ(1-42) red subunit and (**B**) the Aβ(1-42) green subunit that comprise the Aβ(1-42) tetramer.

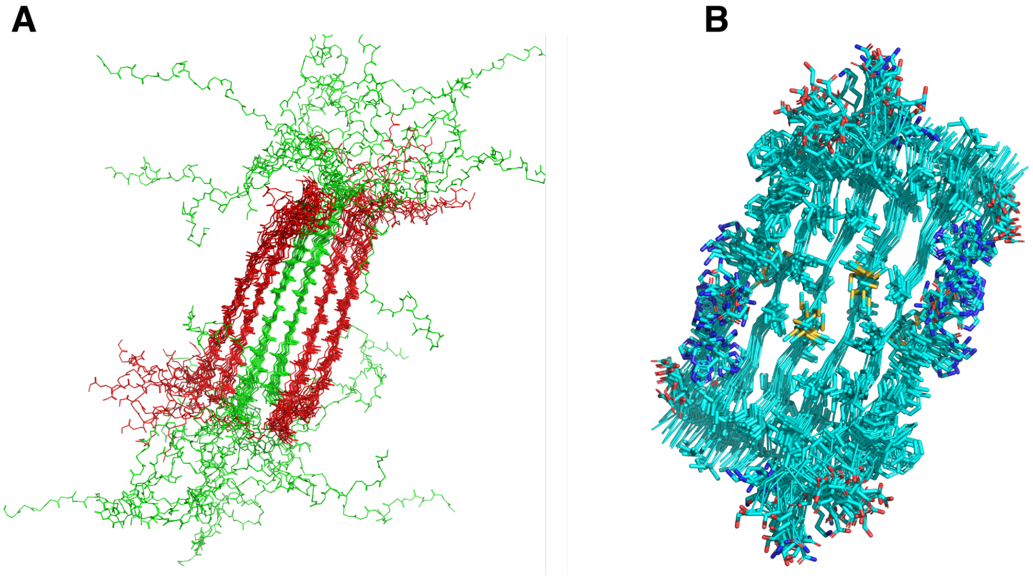


**fig. S7.** NMR ensemble of the 10 lowest energy structures calculated for the Aβ(1-42) tetramer depicting (**A**) all backbone atoms and (**B**) backbone and side chain atoms of the β-sheet core. Residues in the six-stranded β-sheet core are used for superposition.

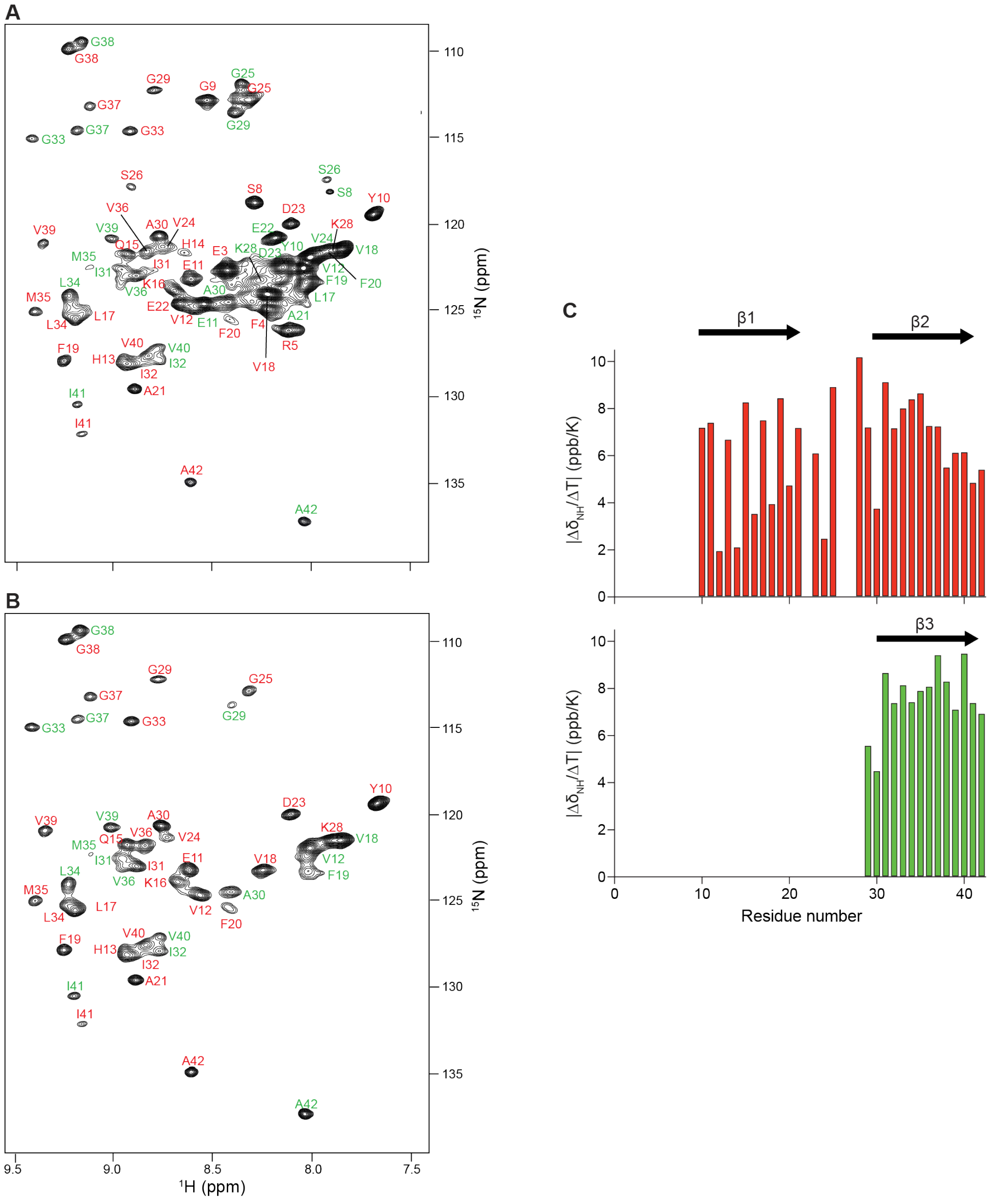


**fig. S8.** Water accessibility of amide protons of the Aβ(1-42) tetramer prepared in DPC. 2D [$^{1}$H,$^{15}$N]-HSQC spectra obtained at (**A**) pH 8.5 and (**B**) pH 9.5. (**C**) Amide proton temperature coefficients versus residue number for the red (top) and the green (bottom) Aβ(1-42) subunits. Secondary structural elements are shown at the top with their corresponding number. Arrows indicate β-strands and helical symbols helices.

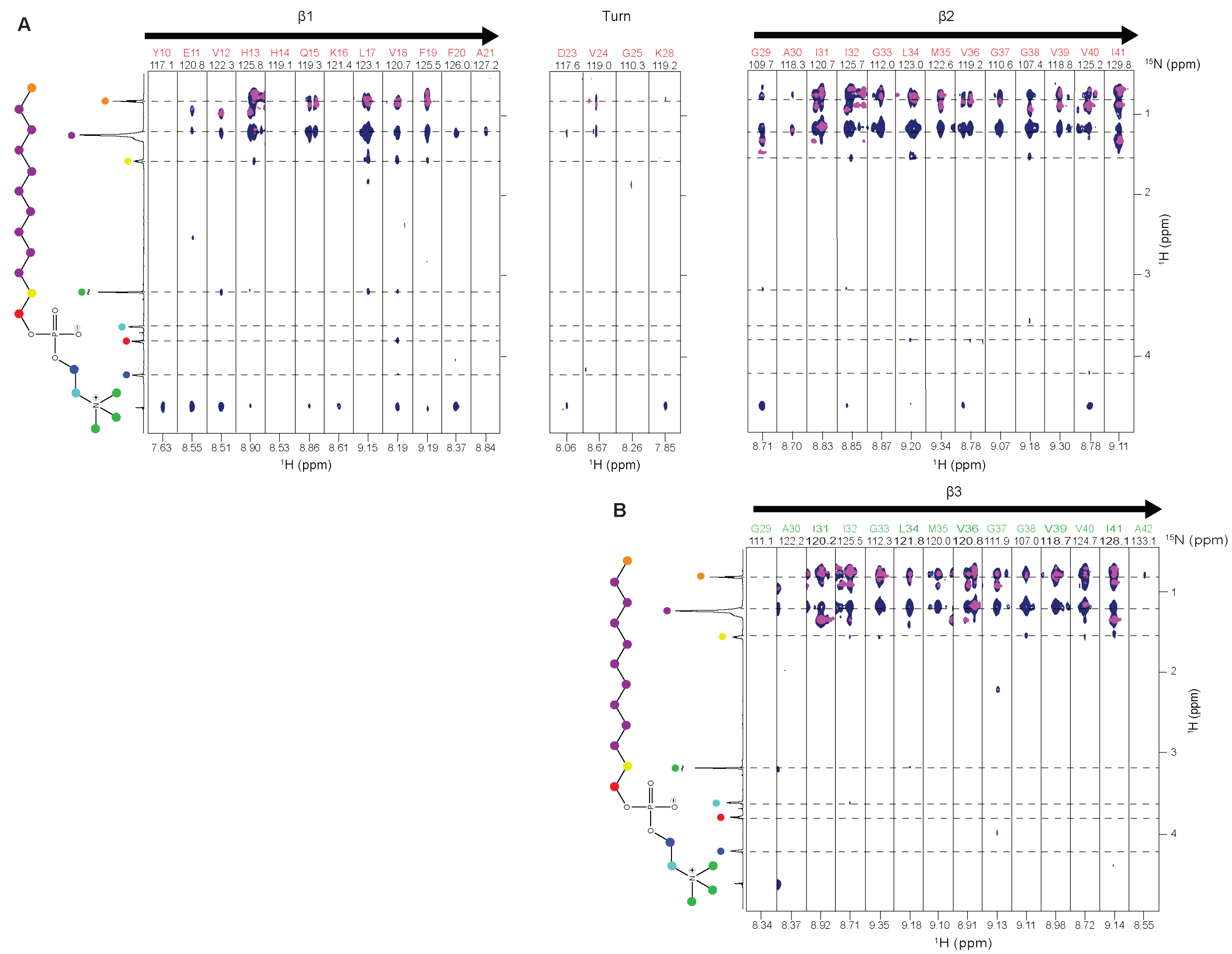


**fig. S9.** Strips of 3D $^{15}$N-resolved [$^{1}$H,$^{1}$H]-NOESY spectrum measured using a uniform $^{2}$H,$^{15}$N, AILV-$^{13}CH_3$ Aβ(1-42) tetramer sample in deuterated DPC micelles (pink) and protonated DPC micelles (blue). The strips were taken at the $^{15}$N chemical shifts of the residues indicated at the top and are centered around the respective amide proton chemical shifts for residues assigned to (**A**) the red Aβ(1-42) subunit and (**B**) the green Aβ(1-42) subunit. On the left side of the strips, one dimensional $^{1}$H NMR spectrum of DPC, measured with the same sample as the 3D $^{15}$N-resolved [$^{1}$H,$^{1}$H]-NOESY and the chemical structure of DPC. The CHn moieties of interest in this study are color-coded with magenta circles to indicate the CHn groups of the hydrophobic tails, and with green circles to identify the polar head methyls of DPC. Above the strips, arrows indicate β-strands.

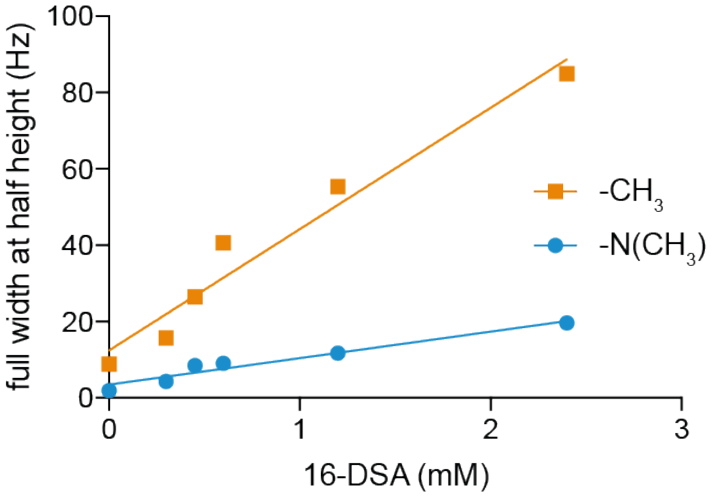


**fig. S10.** 16-DSA is properly inserted into DPC micelles. Line widths $\Delta\nu_{1/2}$ (full width at half weight) of DPC resonance in the 1D $^1$H NMR spectra plotted against the concentration of 16-DSA. The data for the –CH3 resonance at 0.85 ppm (red line) and the –N(CH3)3 group at 3.26 ppm (blue line) are shown. The ε values obtained from linear fits are indicated.

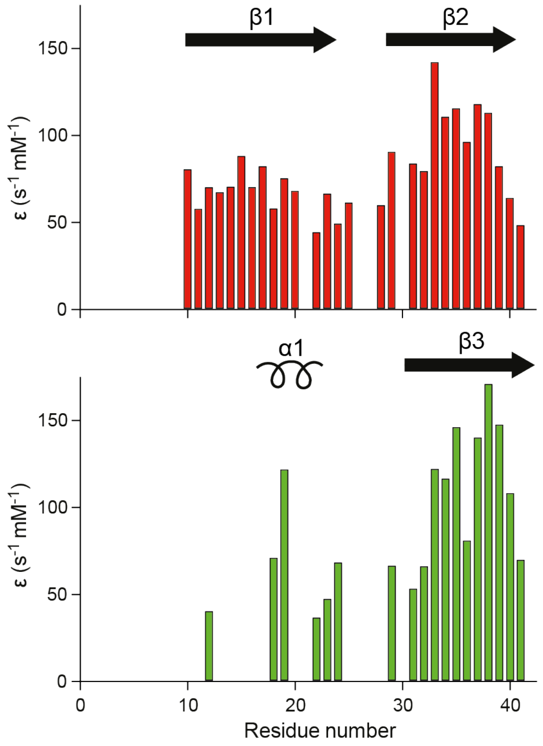


**fig. S11.** Effect of 16-DSA effect on backbone amide protons of the Aβ(1-42) tetramer. Paramagnetic relaxation enhancement, ε, versus residue number for the red (top) and the green (bottom) Aβ(1-42) tetramer subunits estimated from the decay of the resonances in 2D [$^{1}$H,$^{15}$N]-HSQC spectra upon titration with the paramagnetic relaxation enhancement 16-DSA. Secondary structural elements are shown at the top with their corresponding number. Arrows indicate β-strands and helical symbols helices.

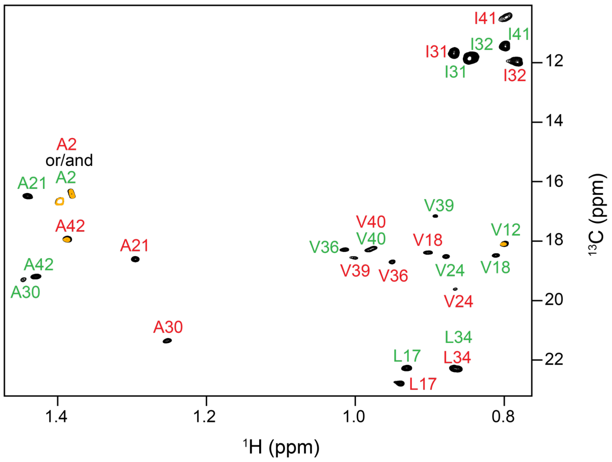


**fig. S12.** DPC micelles are arranged on both sides of the β-sheet core of the Aβ(1-42) tetramer. The 2D [$^{1}$H,$^{13}$C]-TROSY spectrum of Aβ(1-42) tetramer prepared using selectively $^{13}$C methyl-protonated AILV and otherwise uniform $^{2}$H,$^{15}$N Aβ(1-42) in DPC in the absence (black spectra) and in the presence of 0.6 mM 16-DSA (orange spectra). All signals corresponding to methyl groups located at either face of the β-sheet core disappeared while those located in the flexible N-termini remained, suggesting that the two faces of the six-stranded β-sheet core were in contact with the hydrophobic tail of DPC molecules.

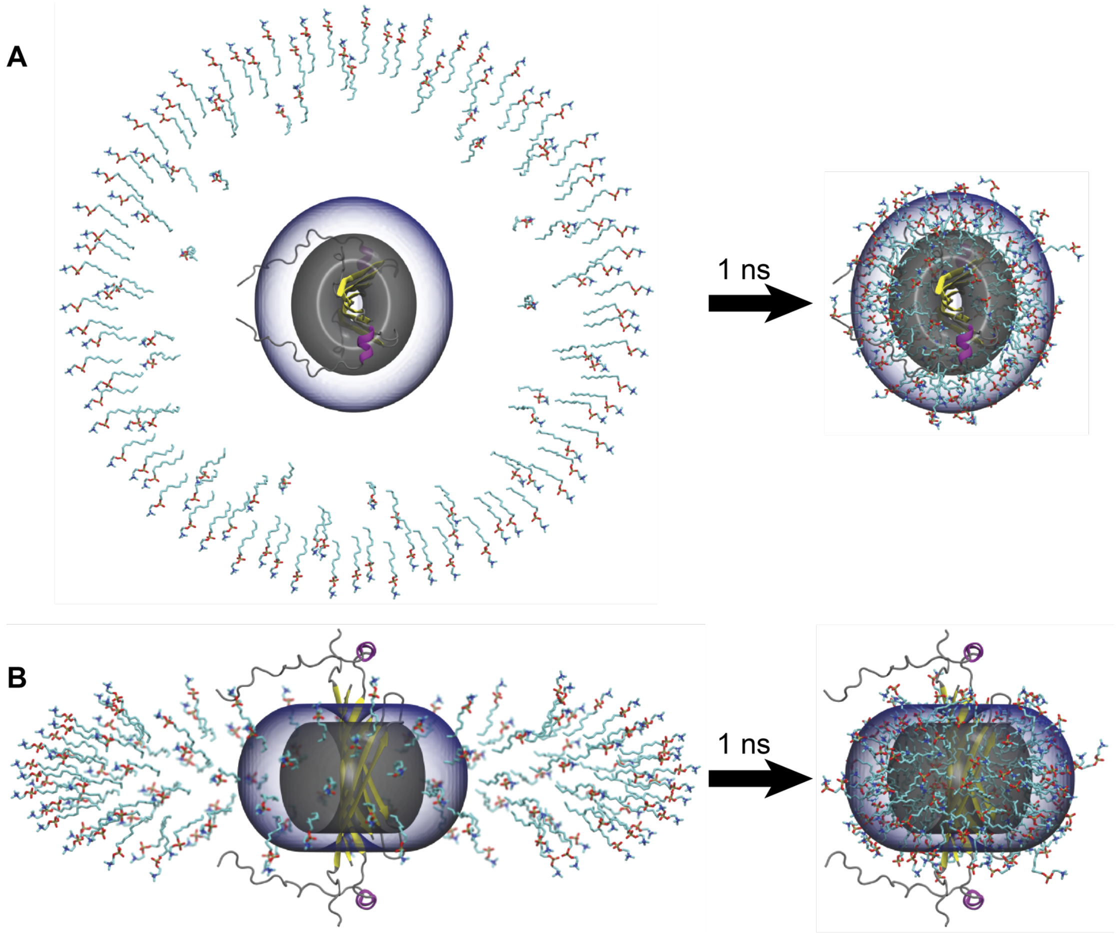


**fig. S13.** SimShape accelerated detergent assembly. Detergent molecules were initially placed far away from the protein assembly into the toroid-shaped potentials and left in an implicit solvent simulation with attractive grid potentials over the course of 1 ns, shown from top **(A)** and side **(B)**.

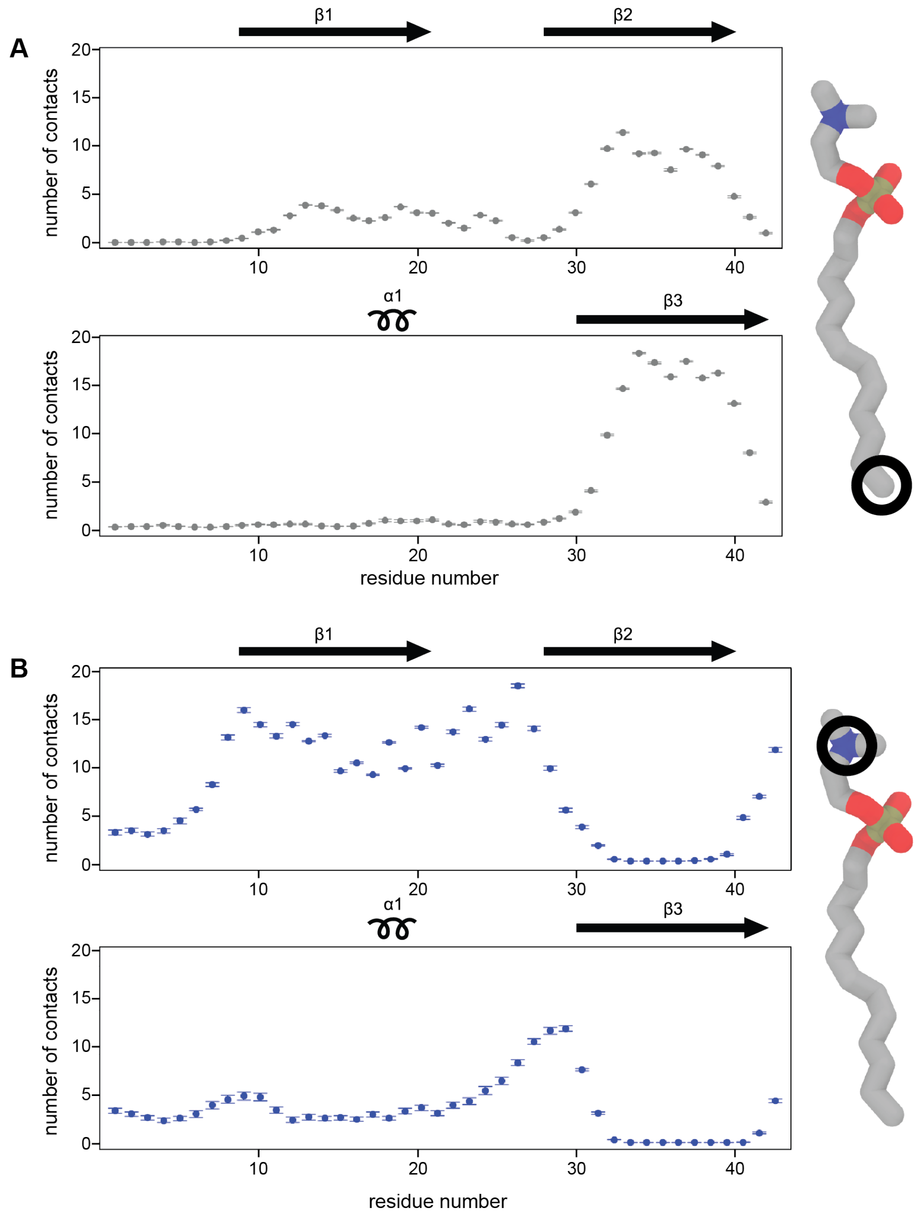


**fig. S14.** Contacts between backbone nitrogen atoms and DPC. The average number of per-residue contacts with **(A)** DPC terminal tail carbon atoms and **(B)** DPC headgroup nitrogen atoms, summed over symmetric chains. Values are reported as the mean over eight independent replicates +/- S.E.M. DPC molecules drawn on the right of each panel show the atom used to define each contact with a black circle. Secondary structural elements are shown at the top of each panel with their corresponding number. Arrows indicate β-strands and helical symbols helices.

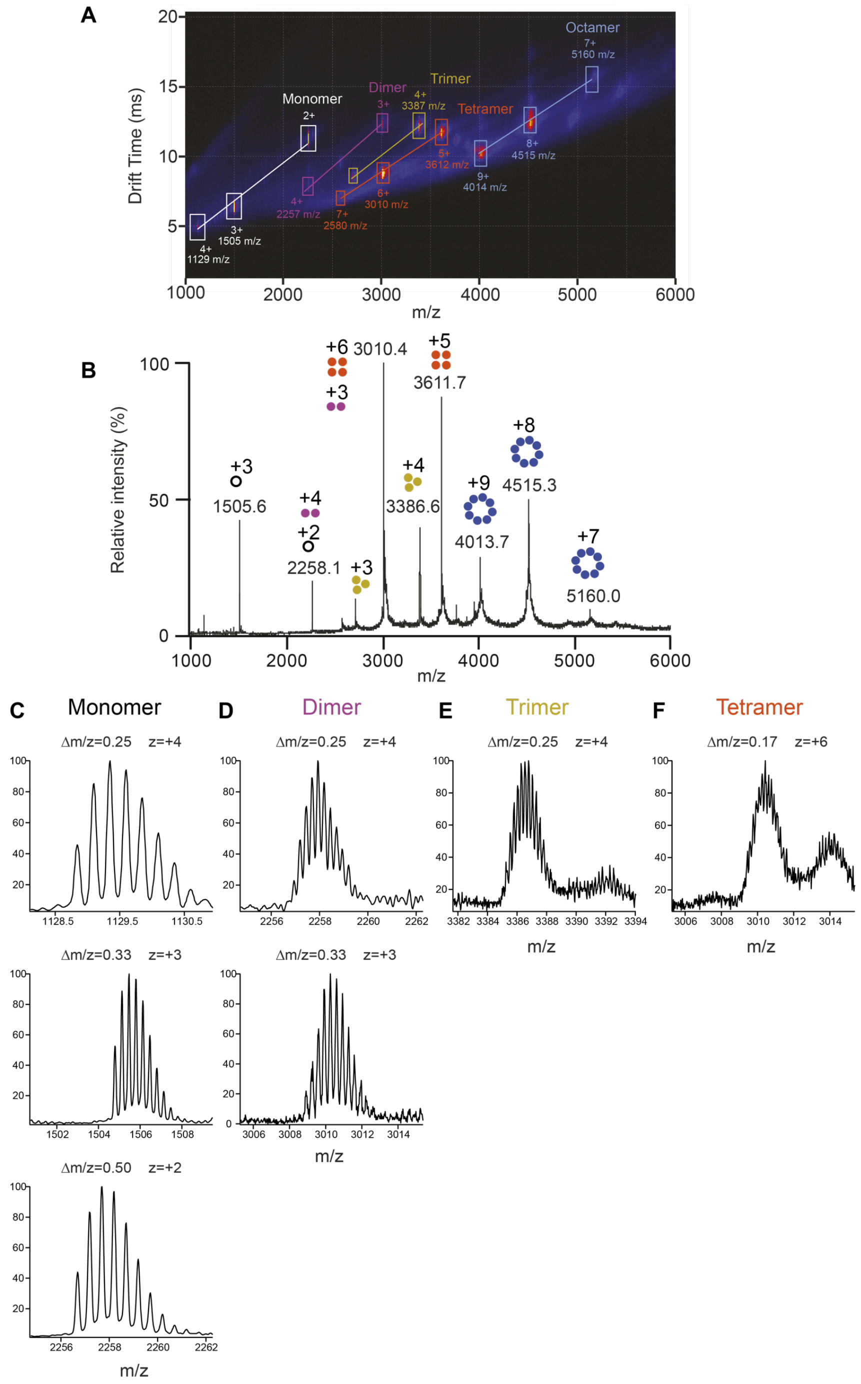


**fig. S15.** ESI-IM-MS analysis of *βPFOs*$_{LOW\_A\beta(1\text{-}42)}$. (**A**) ESI-IM-MS spectrum. Signals corresponding to monomers, dimers, trimers, tetramers, and octamers are indicated, respectively, in white, pink, yellow, orange and blue. The number adjacent to each peak refers to the charge state of the ion. (**B**) Summed m/z spectrum showing the contribution of each oligomeric species to each peak in the spectra. The charge states corresponding to monomers, dimers, trimers, tetramers, and octamers are indicated with schematic drawings and labeled, respectively, in white, pink, yellow, orange and blue. Associated m/z spectrum obtained for mobility peaks associated to (**C**) +4, +3, and +2 charge states of the monomer, (**D**) +4 and +3 charge states of the dimer, (**E**) +4 of the trimer and (**F**) +6 of the tetramer.

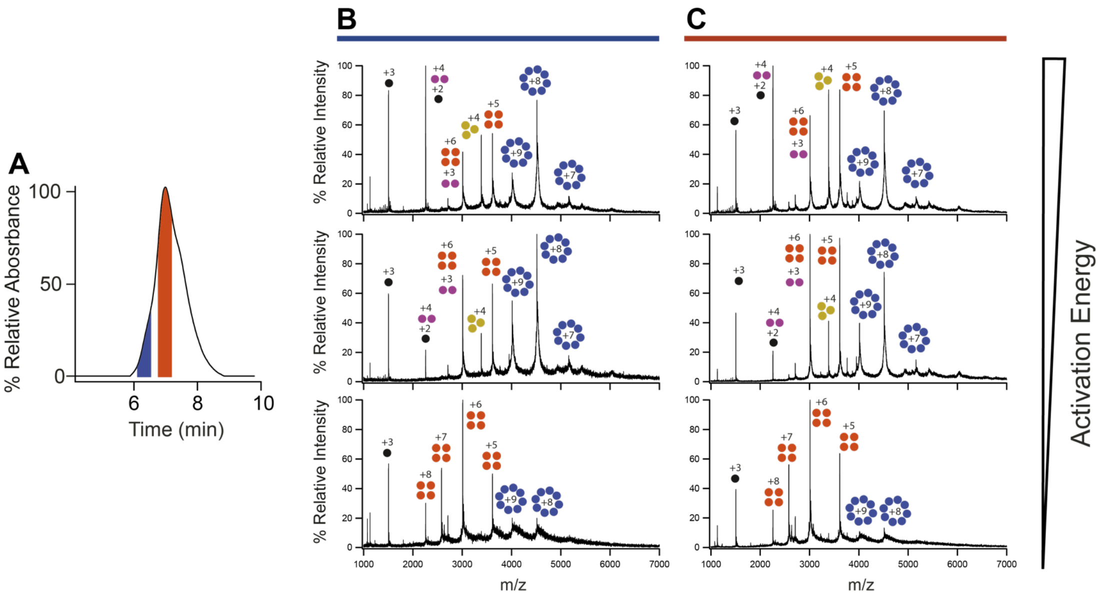


**fig. S16.** Effect of increasing the activation energy of *βPFO*$_{LOW\_A\beta(1\text{-}42)}$ sample by SEC/IM-MS. (**A**) SEC chromatogram obtained with a column equilibrated in C8E5. The mass spectra extracted from the blue and orange SEC peaks and obtained at three activation energies are shown, respectively, with a (**B**) blue and (**C**) orange line on top of them. The charge states corresponding to monomers, dimers, trimers, tetramers, and octamers are indicated with schematic drawings and labeled, respectively, in black, pink, yellow, orange and blue.

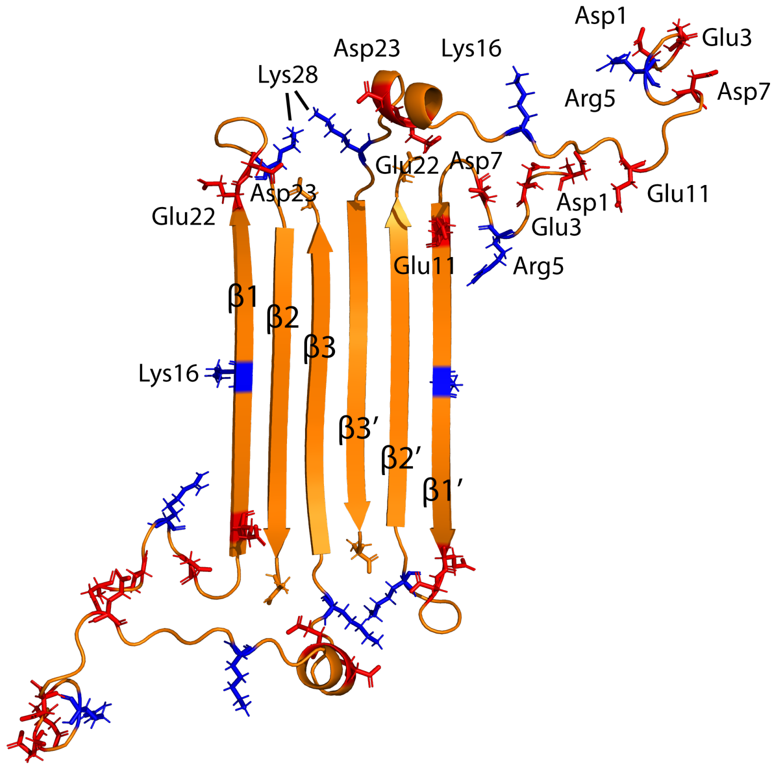


**fig. S17.** 3D structure of the Aβ(1-42) tetramer prepared in DPC with acidic residues highlighted in red (aspartic, glutamic acid, C(t)) and basic residues in blue (N(t) lysine, arginine). The representation shows the proximity of acid and basic residues as potential sites for zero-length cross-linking.

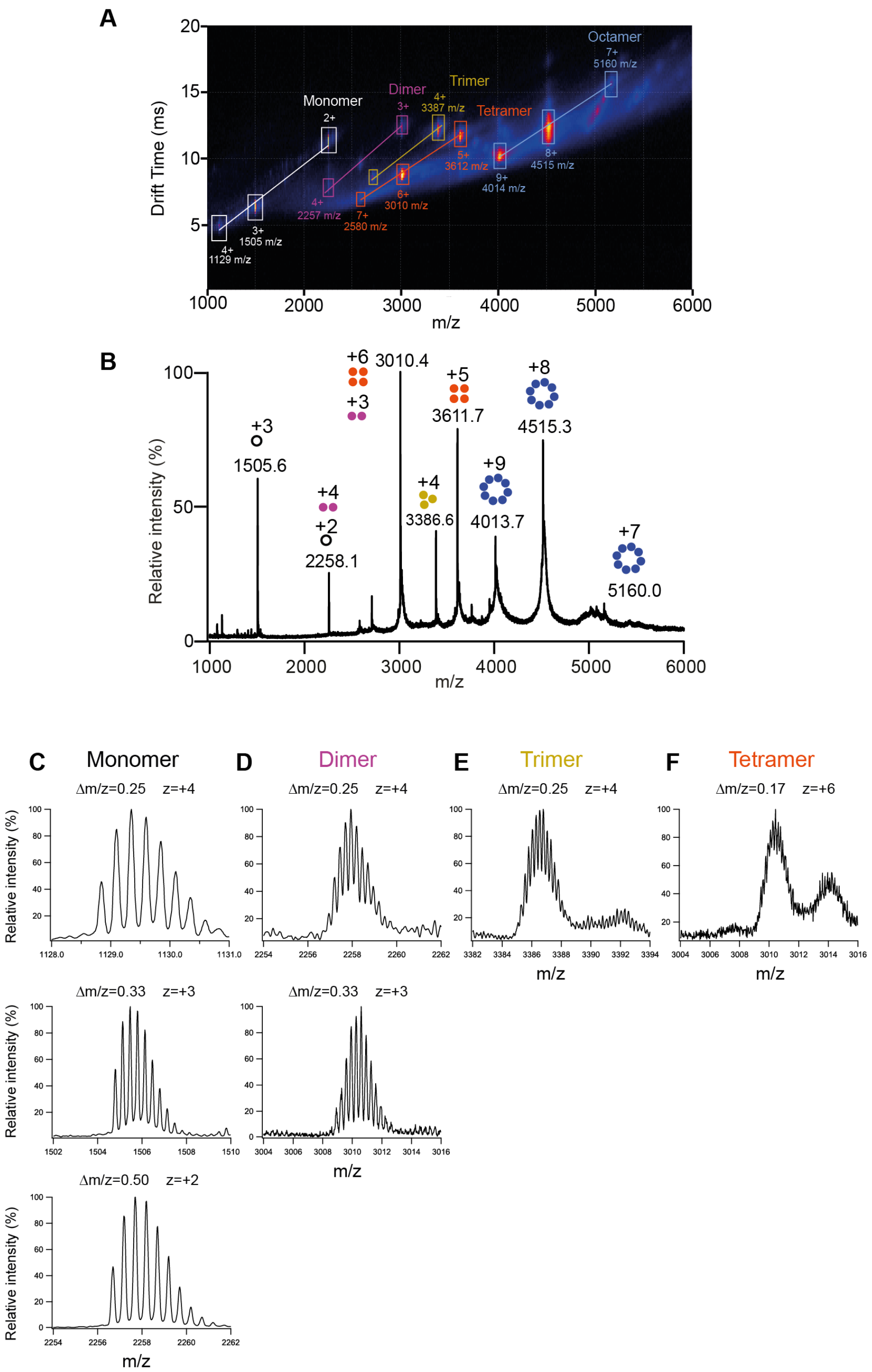


**fig. S18.** ESI-IM-MS analysis of $\beta PFOs_{LOW\_A\beta(1\text{-}42)}$. (**A**) ESI-IM-MS spectrum. Signals corresponding to monomers, dimers, trimers, tetramers, and octamers are indicated, respectively, in white, pink, yellow, orange and blue. The number adjacent to each peak refers to the charge state of the ion. (**B**) Summed m/z spectrum showing the contribution of each oligomeric species to each peak in the spectra. The charge states corresponding to monomers, dimers, trimers, tetramers, and octamers are indicated with schematic drawings and labeled, respectively, in white, pink, yellow, orange and blue. Associated m/z spectrum obtained for mobility peaks associated with (**C**) +4, +3, and +2 charge states of the monomer, (**D**) +4 and +3 charge states of the dimer, (**E**) +4 of the trimer and (**F**) +6 of the tetramer.

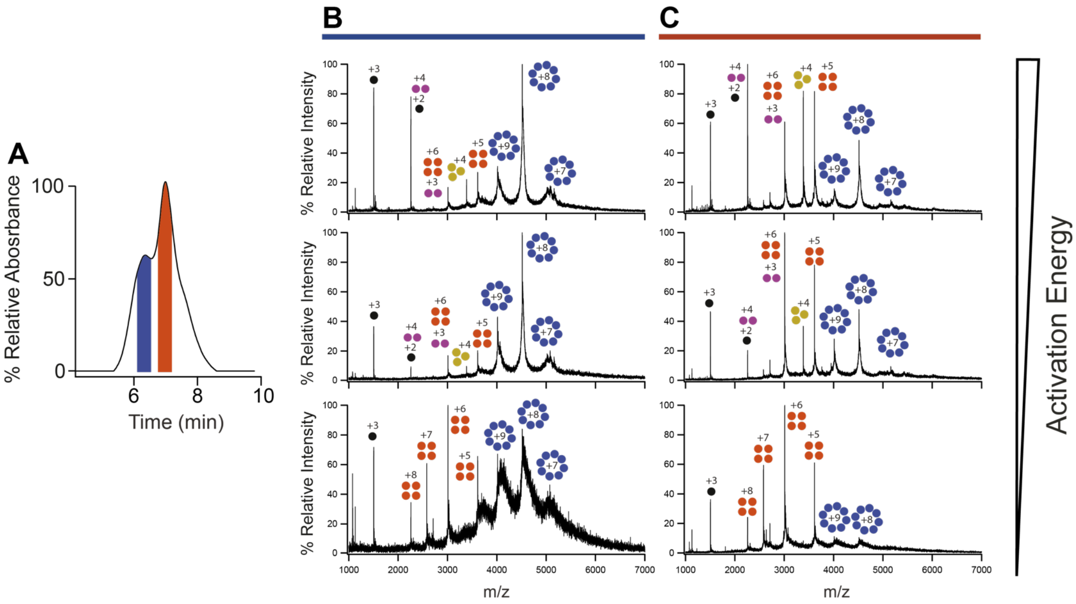


**fig. S19.** Effect of increasing the activation energy of *βPFOs*$_{HIGH\_A\beta(1\text{-}42)}$, by SEC/IM-MS. (**A**) SEC chromatogram obtained with a column equilibrated in C8E5. The mass spectra extracted from the blue and orange SEC peaks and obtained at three activation energies are shown, respectively, with a (**B**) blue and (**C**) orange line on top of them. The charge states corresponding to monomers, dimers, trimers, tetramers, and octamers are indicated with schematic drawings and labeled, respectively, in black, pink, yellow, orange and blue.

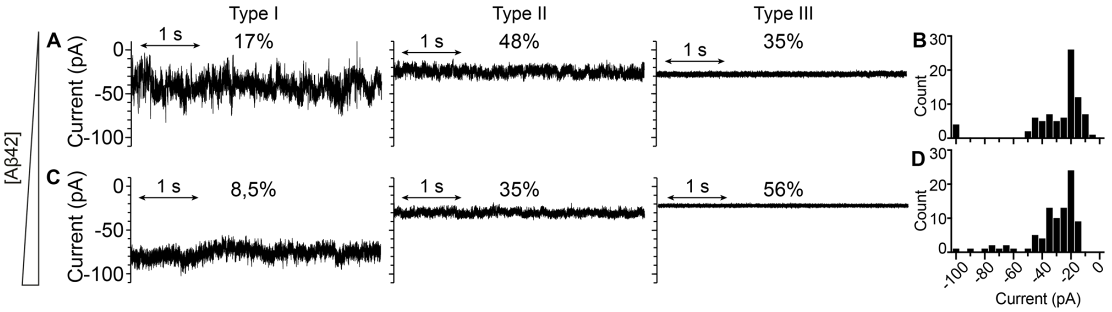


**fig. S20.** *βPFOs*$_{A\beta(1\text{-}42)}$ incorporate into lipid bilayers as pores. Typical current traces and relative abundance (shown on top of the current traces as percentages) for type 1, 2, and 3 pores observed for (**A**) *βPFOs*$_{LOW\_A\beta(1\text{-}42)}$ and (**C**) *βPFOs*$_{HIGH\_A\beta(1\text{-}42)}$. Electrical recordings were carried out on diphytanoyl-sn-glycero-3-phosphocholine planar lipid bilayers. All point histogram for type 2 and type 3 for (**B**) *βPFOs*$_{LOW\_A\beta(1\text{-}42)}$ and (**D**) *βPFOs*$_{HIGH\_A\beta(1\text{-}42)}$ under an applied potential of -100 mV.

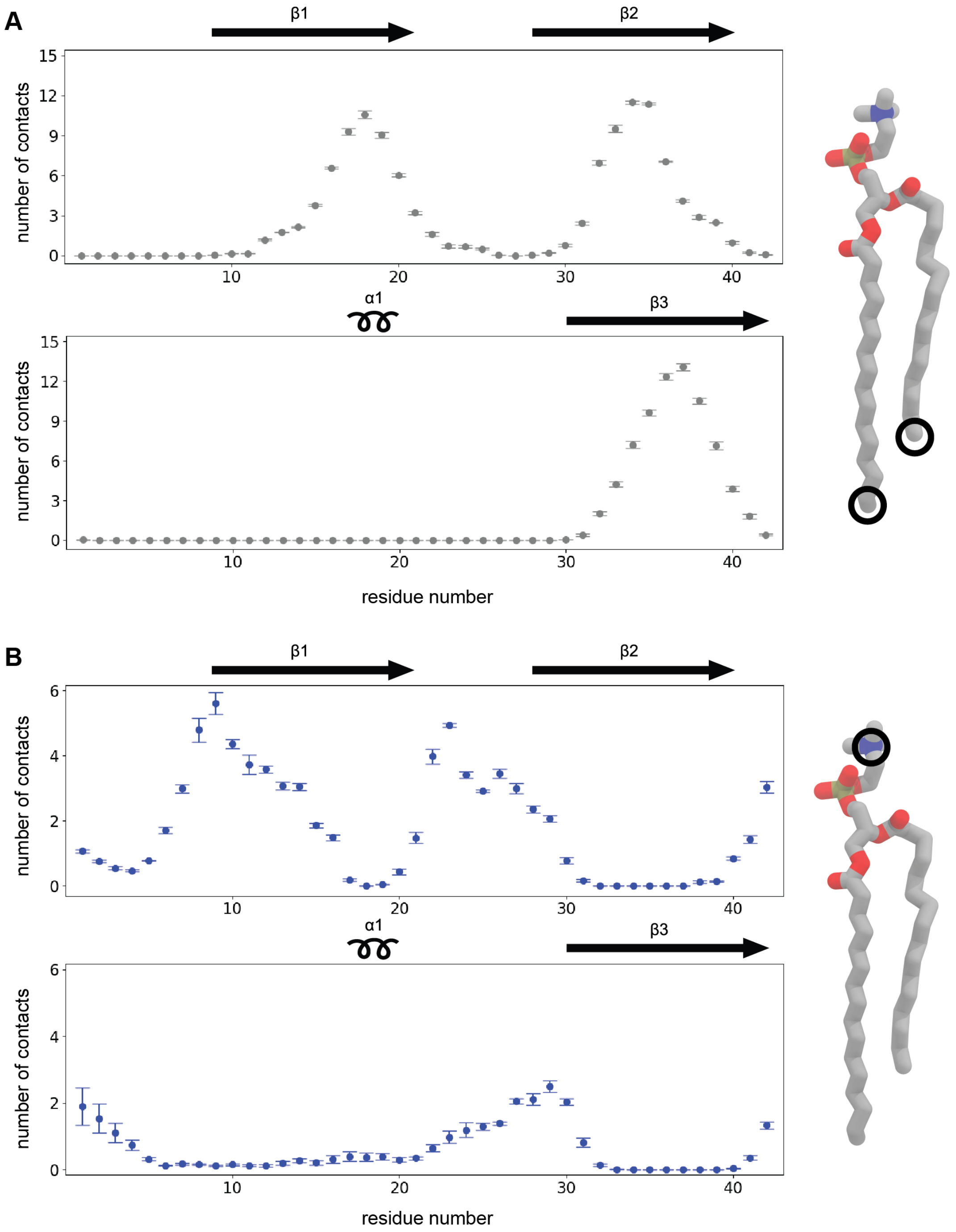


**fig. S21.** Contacts between tetramer backbone nitrogen atoms and DPPC. The average number of per-residue contacts with DPPC hydrophobic tail atoms (A) and DPPC headgroup nitrogen atoms (B), summed over symmetric chains. Values are reported as the mean over three independent replicates +/- S.E.M. DPPC molecules (right) mark the atom used to define each contact with a black circle.

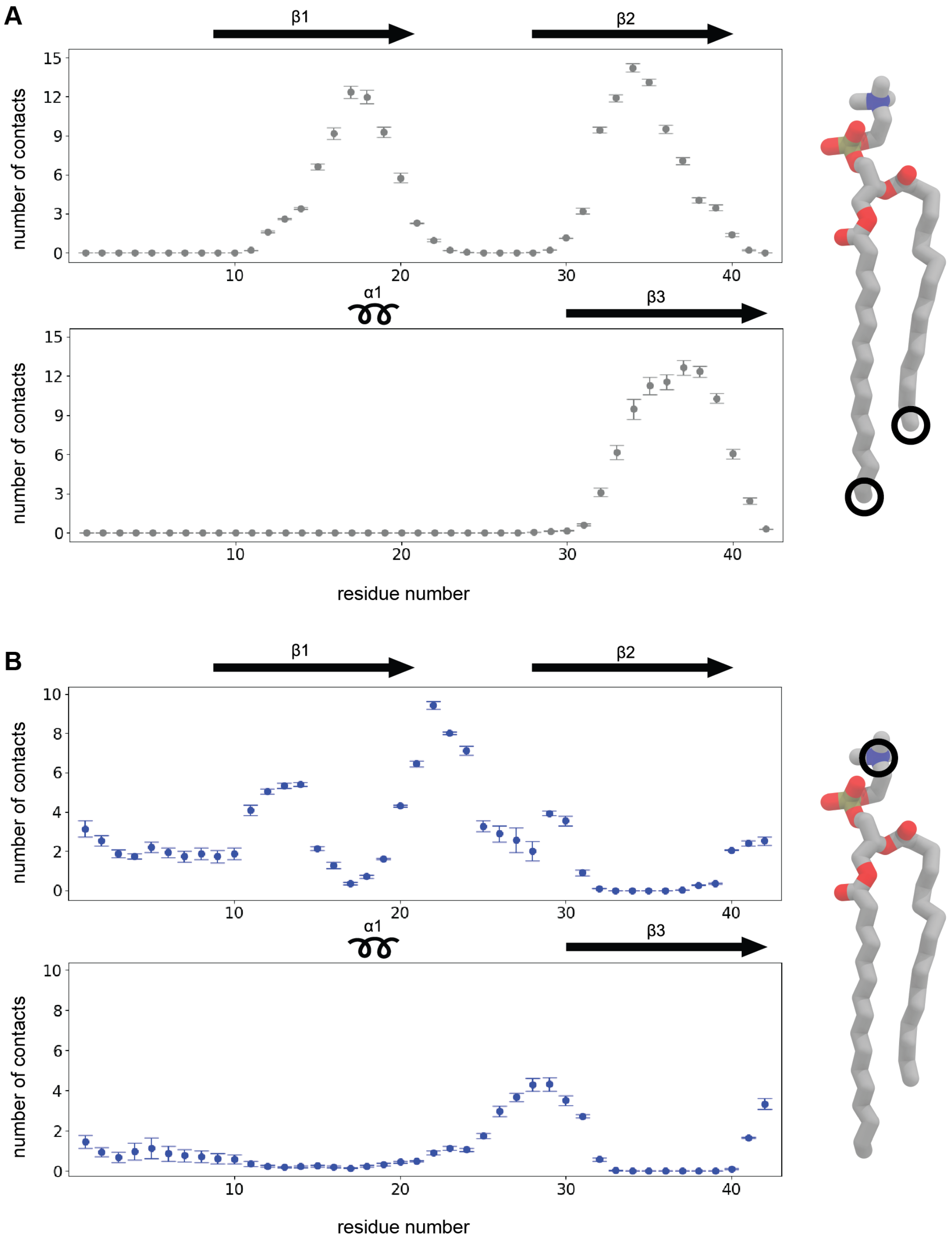


**fig. S22.** Contacts between octamer backbone nitrogen atoms and DPPC. The average number of per-residue contacts with DPPC hydrophobic tail atoms (A) and DPPC headgroup nitrogen atoms (B), summed over symmetric chains. Values are reported as the mean over three independent replicates +/- S.E.M. DPPC molecules (right) mark the atom used to define each contact with a black circle.

### Supporting Tables

***table S1***: Number of unique NOE-derived distance restraints used for structure calculation of Aβ(1-42) tetramers (per dimer unit).

| NOE distance restraints | all | HN-HN | HN-Met | Met-Met |
|---|---|---|---|---|
| total | **157** | 45 | 87 | 25 |
| intra-monomer | **107** | 35 | 60 | 12 |
| intra-residual (\|i-j\| = 0) | **22** | 0 | 22 | 0 |
| sequential (\|i-j\| = 1) | **44** | 24 | 20 | 0 |
| medium-range (2 ≤ \|i-j\| < 5) | **14** | 3 | 5 | 6 |
| long-range (\|i-j\| ≥ 5) | **26** | 8 | 12 | 6 |
| ambiguous | **1** | 0 | 1 | 0 |
| inter-monomer | **30** | 7 | 15 | 8 |
| inter-dimer | **13** | 3 | 5 | 5 |
| ambiguous | **7** | 0 | 7 | 0 |

***table S2***: Structure and restraints statistics for the Aβ(1-42) tetramer ensemble

| **Number of restraints (per dimer)** | |
|---|---|
| NOE distance restraints | |
| Intra-momomer | |
| Intra-residue (\|i-j\| = 0) | 22 |
| Sequential (\|i-j\| = 1) | 44 |
| Medium-range (2 ≤ \|i-j\| < 5) | 14 |
| Long-range (\|i-j\| ≥ 5) | 26 |
| Ambiguous | 1 |
| *Total* | *107* |
| Inter-monomer | 30 |
| Inter-dimer | 13 |
| Ambiguous | 7 |
| *Total* | *157* |
| Dihedral angle restraints (ϕ/ψ) | 82 (41/41) |
| Hydrogen-bond restraints* | |
| Intra-momomer | 24 |
| Inter-momomer | 28 |
| Inter-dimer | 12 |
| *Total* | *64* |

| | |
|---|---|
| **Restraints statistics[a]** | |
| RMS of distance violations | |
| NOE restraints | 0.007 ± 0.002 Å |
| H-bonds restraints | 0.001 ± 0.001 Å |
| Distance violations (count per conformer) | |
| > 0.5 Å | 0 |
| > 0.3 Å | 0 |
| > 0.1 Å | 0 |
| RMS of dihedral violations | 0.33 ± 0.11° |
| Dihedral violations > 5° (count per conformer) | 0 |
| **RMS from idealized covalent geometry** | |
| bonds | 0.003 ± 0.001 Å |
| angles | 0.376 ± 0.007° |
| impropers | 0.898 ± 0.091° |
| **Structural quality[a]** | |
| Ramachandran statistics[b] | |
| Most favoured regions | 76.0 ± 4.0 % |
| Allowed regions | 23.7 ± 3.7 % |
| Disallowed regions | 0.3 ± 0.6 % |
| Molprobity | |
| Clashscore | 1.88 ± 0.64 (99th percentile) |
| Coordinates precision[c] | |
| Backbone atoms (core β-sheet[d], tetramer) | 0.77 ± 0.27 Å |
| Heavy atoms (core β-sheet, tetramer) | 1.34 ± 0.18 Å |
| Backbone atoms (core β-sheet, dimer) | 0.54 ± 0.14 Å |
| Heavy atoms (core β-sheet, dimer) | 1.19 ± 0.12 Å |

*One hydrogen-bond is encoded with 2 restraints (HN…O and N…O)
[a] Average values and standard deviations over the 15 models
[b] Percentage of residues in the Ramachandran plot regions determined by PROCHECK (Laskowski et al., 1993)
[c] Average root mean square deviation (RMSD) over the 15 conformers with respect to the average structure.
[d] Residues β1-β2-β3-β1'-β2'-β3' (only β1-β2-β3 for dimer)

***table S3***: Acquisition parameters of the NMR experiments carried out throughout this study

| Experiment | D1 | NS | NUS % | F3<br>AQ (ms) / SW (ppm) | F2<br>AQ (ms) / SW (ppm) | F1<br>AQ (ms) / SW (ppm) | NOE mix (s) | Exp time (h) |
|---|---|---|---|---|---|---|---|---|
| 3D HNCA (pH 8.5) | 1.5 | 8 | 25 | 71/18 | 40/32 | 25/46 | | 93 |
| 3D HNCA (pH 9.5) | 1.5 | 8 | 30 | 71/18 | 40/32 | 45/26 | | 89 |
| 3D HNCACB (pH 8.5) | 1.5 | 16 | 30 | 71/18 | 18/46 | 14/75 | | 129 |
| 3D HNCACB (pH 9.5) | 1.2 | 16 | 30 | 71/18 | 22/26 | 12/75 | | 64 |
| HNCO (pH 9.5) | 1.5 | 64 | 50 | | 63/40 | 79/47 | | 18 |
| 3D HN-NH NOESY | 0.25 | 16 | 25 | 70/6 | 30/3 | 37/26 | 0.08 | 94 |
| 3D (Hme)Cme([C] CA)CO | 1.0 | 8 | 50 | 61/14 | 14/16 | 10/14 | | 9 |
| 3D (Hme)Cme([C] CA)NH | 1.0 | 16 | 50 | 61/14 | 22/25 | 11/16 | | 22 |
| 3D Hme(Cme[C]CA) NH | 1.0 | 16 | 50 | 61/14 | 22/25 | 18/2 | | 17.5 |
| 3D (H)C-TOCSY-C-TOCSY-(C)H | 1.25 | 8 | 30 | 57/20 | 8.6/66 | 11/16 | | 19.5 |
| 3D Hm-CmHm NO-ESY | 0.2 | 16 | 50 | 57/10 | 15/14 | 36/1 | 0.3 | 8.5 |
| 3D Cm- CmHm NO-ESY | 0.3 | 64 | 65.1 | 57/10 | 15/14 | 15/14 | 0.2 | 66.5 |
| 3D Hm-NH NOESY | 0.2 | 16 | 50 | 57/20 | 27/26 | 36/1 | 0.3 | 12.5 |
| 3D Cm-NH NOESY | 0.2 | 16 | 50 | 57/20 | 27/26 | 15/14 | 0.3 | 17 |

***table S4***: Comparison of the reported CCS' for the calibrants used to calibrate IMS with the CCS' obtained for the tetramer structures and octamer models using the projection approximation algorithm within the IMPACT software. The octamer models are based on the association of two Aβ(1-42) tetramers to form a β-barrel or a β-sandwich structure.

| **Protein** | **MW (Da)** | **CCS Ω (Å²)** | **Z (charge)** | **CCS' 'Ω (Å²)** |
|---|---|---|---|---|
| Tetramer w/o flex. | 9652 | 1270±4 | 7 | 417 |
| *Cytochrome C* | 12000 | 1240 | 6 | 475 |
| Octamer β-sandwich w/o flex. | 19304 | 1918±6 | 9 | 490 |
| *β-lactoglobulin monomer* | 18000 | 1660 | 7 | 545 |
| Octamer β-barrel w/o flex. | 19304 | 2322±3 | 9 | 593 |
| *β-lactoglobulin dimer* | 36000 | 2850 | 11 | 596 |
| *Concanavalin A* | 103000 | 5550 | 20 | 638 |
| *Alcohol dehydrogenase* | 143000 | 6940 | 24 | 665 |
| *Glutamate dehydrogenase* | 336 | 12800 | 38 | 775 |
| *Pyruvate kinase* | 237000 | 10300 | 30 | 790 |
| Tetramer | 18056 | 2545±8 | 7 | 836 |
| Octamer β-sandwich | 36112 | 4173±11 | 9 | 1066 |
| Octamer β-barrel | 19304 | 4348±15 | 9 | 1111 |

### Supporting References


1. M. Serra-Batiste *et al.*, *Curr Chem Biol.* **11**, 50–62 (2017).

2. R. Kerfah *et al.*, *J. Biomol. NMR*. **61**, 73–82 (2015).

3. P. K. Mandal, J. W. Pettegrew, *Neurochem Res*. **29**, 1–6 (2004).

4. C. R. Sanders, F. Sönnichsen, *Magn. Reson. Chem.* **44**, S24–S40 (2006).

5. P. Rossi, Y. Xia, N. Khanra, G. Veglia, C. G. Kalodimos, *J. Biomol. NMR*. **66**, 259–271 (2016).

6. M. Mayzel, K. Kazimierczuk, V. Y. Orekhov, *Chem. Commun.* **50**, 8947–8950 (2014).

7. V. Y. Orekhov, V. A. Jaravine, *Prog. Nucl. Magn. Reson. Spectrosc.* **59**, 271–292 (2011).

8. H. Shao, S. Jao, K. Ma, M. G. Zagorski, *J. Mol. Biol.* **285**, 755–773 (1999).

9. T. Cierpicki, J. Otlewski, *J. Biomol. NMR*. **21**, 249–261 (2001).

10. M. Serra-Batiste *et al.*, *Proc. Natl. Acad. Sci. U.S.A.* **113**, 10866–10871 (2016).

11. M. F. Bush *et al.*, *Anal. Chem.* **82**, 9557–9565 (2010).

12. T. M. Allison, M. Landreh, J. L. P. Benesch, C. V. Robinson, *Anal. Chem.* **88**, 5879–5884 (2016).

13. A. Leitner *et al.*, *Proc. Natl. Acad. Sci. USA*. **111**, 9455–9460 (2014).

14. M. Cadene, B. T. Chait, *Anal. Chem.* **72**, 5655–5658 (2000).
